# Lipotoxic fingerprints in clinically relevant postoperative pancreatic fistula: fatty acid–driven cytotoxicity targets cells involved in anastomotic healing

**DOI:** 10.64898/2026.02.12.705517

**Authors:** Johannes D. Lettner, Marvin Schwarzer, Simon Lagies, Bernd Kammerer, Stephanie Mewes, Sophia Chikhladze, Stefan Fichtner-Feigl, Geoffroy Andrieux, Dietrich A. Ruess, Uwe A. Wittel

**Author notes:** Corresponding author, Johannes D. Lettner, Hugstetterstraße 55, 79106 Freiburg Germany, mail, Tel.: +49 0761/270 23650. Equal Contribution, shared last Authorship (Dietrich A. Ruess, Uwe A. Wittel).

## Abstract

**Background & Aims:** Clinically relevant postoperative pancreatic fistula (CR-POPF) remains a major cause of morbidity following pancreatic surgery, potentially due to fatty acid release by lipase activity. This study investigated how the biochemical composition of CR-POPF effluents drives cellular injury and transcriptional stress responses.

**Methods:** Drain effluents from 14 patients undergoing pancreatoduodenectomy (7 with CR-POPF and 7 controls) were analyzed using gas chromatography–mass spectrometry. Candidate lipids were tested on human foreskin fibroblasts, mesothelial cells, and pancreatic epithelial cells using viability and cytotoxicity assays. Effluents were applied directly to cultures, and RNA sequencing was performed on cells exposed to the two most cytotoxic CR-POPF samples.

**Results:** Metabolomic profiling revealed lipolytic traits characterized by long-chain saturated fatty acids, including palmitic and stearic acid, and the palmitic acid monoacylglycerol monopalmitin, in drain effluents. These fatty acids accounted for over 70% of the variance in multivariate metabolomic analyses between CR-POPF and control groups. Dose–response assays confirmed concentration-dependent cytotoxicity (p < 0.0001), with a subtoxic threshold of 0.2 mM. Two effluents (AES1448 and GR1479) consistently reduced cell viability across models (F > 19, p < 0.0001). Transcriptomic profiling showed enrichment of inflammatory, unfolded-protein, and stress-response pathways, along with suppression of proliferation modules. GR1479 induced metabolic adaptation, whereas AES1448 and monopalmitin triggered overt lipotoxic stress.

**Conclusions:** Lipolysis-derived lipids may mediate stromal and mesothelial injury in CR-POPF. Integrating metabolomic, functional, and transcriptomic data uncovers a spectrum of cellular responses, spanning from adaptive remodeling to lipolysis-driven proteotoxic stress. These findings support lipid toxicity as a biochemical property of CR-POPF and a potential target for prevention.

**Synopsis:** This study identifies long-chain saturated fatty acids in postoperative pancreatic effluents as key mediators of cytotoxic and inflammatory stress. Integrating metabolomic and transcriptomic analyses link effluent composition directly to cellular injury and impaired healing after pancreatic surgery.

Graphical abstract

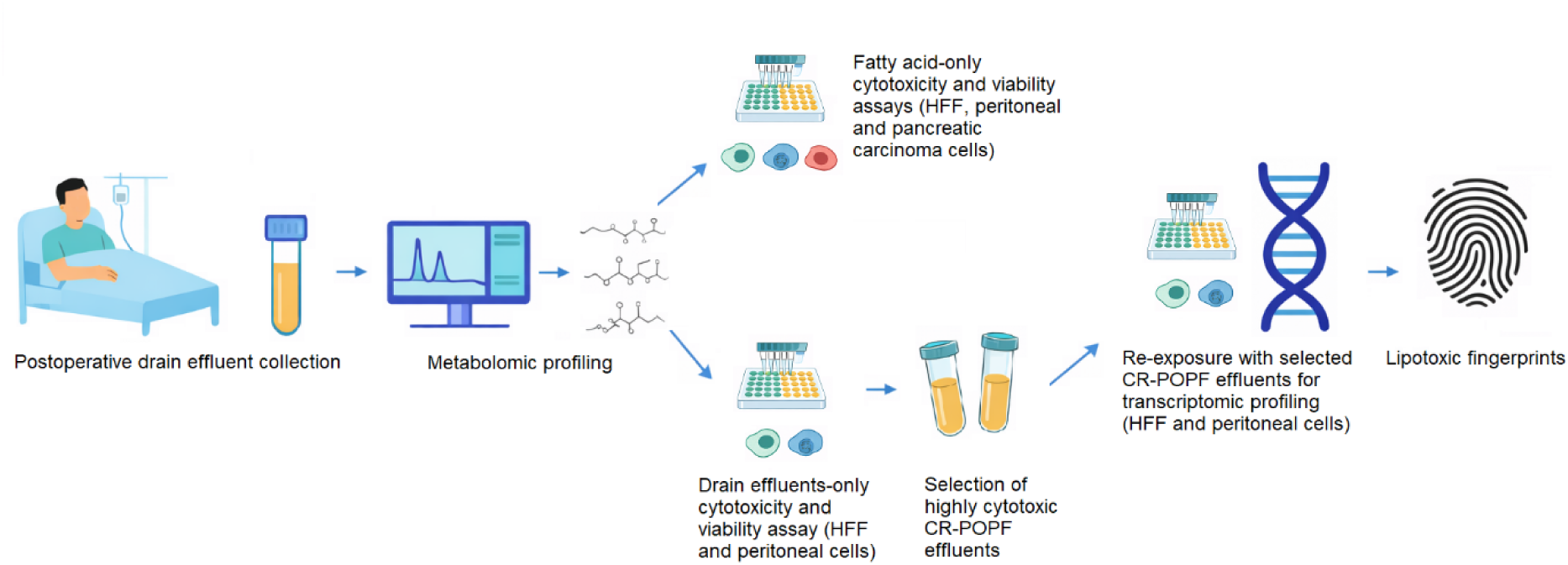

## Introduction

Clinically relevant postoperative pancreatic fistula (CR-POPF) remains one of the most serious complications after pancreatic surgery. Despite major advances in surgical techniques and perioperative care, its incidence remains at up to 15%, continuing to drive postoperative morbidity and mortality (1–3). The underlying mechanisms, however, remain incompletely understood (4–6). Beside established mechanical and anatomical risk factors such as the texture of the soft glands, the size of the ducts, and the body mass index, recent studies suggest that locally generated free fatty acids may represent an additional biochemical factor contributing to the development of clinically relevant postoperative pancreatic fistula (5,7). This is because leakage of pancreatic juice into the peritoneal cavity promotes lipase-mediated intraperitoneal lipolysis, resulting in the accumulation of free fatty acids, which can be measured in postoperative drain fluid (7). Experimental studies have shown that saturated free fatty acids such as palmitic acid can have a direct cytotoxic effect on pancreatic cells by inducing oxidative stress, endoplasmic reticulum stress, and pro-inflammatory signaling pathways (8). This mechanism resembles the pathophysiology of acute pancreatitis, in which lipase-mediated degradation of visceral fat leads to the generation of free fatty acids that markedly amplify inflammation and parenchymal necrosis (9).

In line with this concept, the International Study Group of Pancreatic Surgery (ISGPS) has identified postoperative pancreatitis as a key component of CR-POPF pathogenesis (10). Notably, more than half of patients with pancreatic ductal adenocarcinoma (PDAC) exhibit preoperative pancreatic steatosis, suggesting a lipid-rich pancreatic microenvironment in a substantial proportion of surgical patients (11). Furthermore, in obese mice, a lipase-mediated lipotoxicity can convert mild pancreatitis into severe, often lethal disease, while lean animals are protected, highlighting the modifying role of adiposity (12). Importantly, lipolysis of peripancreatic adipose tissue does not result in a uniform lipid milieu but generates defined cytotoxic lipid species (7). Among these, long-chain saturated fatty acids, particularly palmitic acid, are produced during lipase-mediated breakdown of peripancreatic adipose tissue and from circulating triglycerides under hyperlipidemic conditions (13). Beyond its direct lipotoxic effects on pancreatic cells, palmitic acid has been shown to facilitate inflammatory cell death by enabling inflammasome priming and activation, within a priming context that involves NF-κB signaling (14). In parallel, free fatty acids may exert indirect cytotoxic effects via macrophage activation. Palmitic acid induces polarization of macrophages toward a pro-inflammatory M1 phenotype, stimulating the release of cathepsin S-containing exosomes. Uptake of these exosomes by pancreatic acinar cells triggers pyroptotic cell death, thereby aggravating pancreatic inflammation and tissue injury (15). From a clinical perspective, hyperlipidemia is therefore not only of interest as a recognized risk factor for acute pancreatitis, but also as a potential mechanistic driver of lipotoxic inflammation, metabolic disorders, activation of the innate immune system, and death of acinar cells within a unified pathogenic framework (13,15). Nevertheless, direct insights into lipid-driven transcriptional stress responses in stromal compartments relevant for anastomotic healing remain limited. Importantly, CR-POPF is not primarily a consequence of pancreatic acinar cell damage but rather reflects a failure of anastomotic healing. Fibroblasts and peritoneal mesothelial cells are key regulators of extracellular matrix deposition, scar formation, and local immune modulation at the pancreatic anastomosis. Lipotoxic injury affecting these stromal cell populations may therefore directly impair tissue repair, compromise anastomotic integrity, and promote fistula persistence and severity. Accordingly, this study investigates the role of lipolysis in CR-POPF pathogenesis with a specific focus on the transcriptional and cytotoxic responses to lipotoxic stress in fibroblast and peritoneal cell models that are directly involved in postoperative healing.

## Results

### Characteristics of the study cohort

Samples of a total of 14 patients, seven with clinically relevant postoperative pancreatic fistula (CR-POPF) and seven without (non-POPF) were included. There were no significant differences in baseline demographic and preoperative variables between the groups, including age, sex, BMI, ASA classification, diabetes, and duct width (all p > 0.05). Preoperative laboratory values, including serum amylase, lipase and bilirubin, were also similar. Surgical approaches were evenly distributed (50% open, 43% laparoscopic and 7% robotic; p = 0.57) and pancreatic texture was soft in 71% of cases, with no significant differences between groups. As expected, based on the ISGPS definition, persistently elevated drain amylase levels were observed in patients classified as CR-POPF. In contrast, serum amylase showed only a transient increase on postoperative day 2 (p = 0.007), which may reflect local postoperative inflammation rather than established pancreatitis and could be associated with CR-POPF development. Histopathological entities were balanced between groups (p = 0.77). Overall complication rates (63%) and severe complications (Clavien–Dindo ≥ III: 43%) did not differ significantly (Table 1).

**Table 1.** Characteristics of the study cohort. Demographic, surgical, pre-, intra- and postoperative outcomes. Abbreviations: ASA, American Society of Anesthesiologists; BMI, body mass index; IQR, interquartile range; POD, postoperative day; PD, pancreatoduodenectomy; PDAC, pancreatic ductal adenocarcinoma; IPMN, intraductal papillary mucinous neoplasm; AMPAC, ampullary carcinoma; HG, high-grade; LG, low-grade.

|  | All n = 14 | CR-POPF = 7 | Non-POPF = 7 |  |
| --- | --- | --- | --- | --- |
| **Demographic factors** |  |  |  |  |
| Age, years, median [IQR] | 67.5 [56.8-76] | 67 [56-76] | 68 [57-79] | 0.620 |
| Sex, female, n [%] | 4 [28.6%] | 4 [57.1] | 0 [0] | 0.070 |
| BMI, kg/m<sup>2</sup>, median [IQR] | 23.7 [22.2-26.4] | 23 [21.-30.7] | 24.4 [22.-25] | 0.805 |
| **Pre- and intraoperative variables** |  |  |  |  |
| ASA ≥ III, n [%] | 8 [57.1] | 5 [71.4] | 3 [42.9] | 0.405 |
| Diabetes mellitus, yes, n [%] | 6 [42.9] | 2 [28.6] | 4 [57.1] | 0.592 |
| History of pancreatitis, yes, n [%] | 2 [14.3] | 2 [28.6] | 0 [0] | 0.301 |
| Serum Amylase, U/L, median [IQR] | 18.5 [11.5-35.3] | 33 [18-36] | 12 [6-27] | 0.530 |
| Serum Lipase, U/L, median [IQR] | 58 [16.8-105] | 75 [17-129] | 53 [16-67] | 0.620 |
| Serum bilirubin, mg/dl, median [IQR] | 0.6 [0.3-4.8] | 0.4 [0.3-0.6] | 0.9 [0.5-9.3] | 0.073 |
| Procedure |  |  |  | 0.565 |
| Open PD, n [%] | 7 [50] | 4 [57.1] | 3 [42.9] |  |
| Lap PD, n [%] | 6 [42.9] | 3 [42.9] | 3 [42.9] |  |
| Robotic PD, n [%] | 1 [7.1] | 0 [0] | 1 [14.3] |  |
| Duct width, mm, median [IQR] | 3 [3-4] | 3 [3-4] | 4 [3-7] | 0.318 |
| Pancreatic texture, soft, n [%] | 10 [71.4] | 6 [85.7] | 4 [57.1] | 0.559 |
| **Postoperative Outcomes** |  |  |  |  |
| Serum Amylase POD1, U/L, median [IQR] | 39.5 [6-55.5] | 40 [38-72] | 6 [3-42] | 0.165 |
| Serum Amylase POD2, U/L, median [IQR] | 29.5 [5.8-73] | 70 [28-138] | 6 [2-31] | **0.007** |
| Serum Amylase POD3, U/L, median [IQR] | 18 [6.3-68.8] | 65 [24-98] | 7 [2-13] | **0.001** |
| Drain Amylase POD1, U/L, median [IQR] | 248 [18.8-1900] | 1792 [1115-5084] | 24 [3-36] | **0.001** |
| Drain Amylase POD2, U/L, median [IQR] | 669 [16-3952] | 3046 [1287-6935] | 17 [13-104] | **0.001** |
| Drain Amylase POD3, U/L, median [IQR] | 566.5 [11-2121.3] | 1947 [1056-4440] | 11 [4-47] | **0.001** |
| Fistula Grade |  |  |  | **<0.001** |
| B, n [%] | 6 [42.9] | 6 [85.7] | 0 [0] |  |
| C, n [%] | 1 [7.1] | 1 [14.3] | 0 [0] |  |
| Histopathology |  |  |  | 0.771 |
| PDAC Head, n [%] | 5 [35.7] | 2 [28.6] | 3 [42.9] |  |
| AMPAC, n [%] | 4 [28.6] | 2 [28.6] | 2 [28.6] |  |
| HG IPMN, n [%] | 2 [14.3] | 1 [14.3] | 1 [14.3] |  |
| LG IPMN, n [%] | 2 [14.3] | 1 [14.3] | 1 [14.3] |  |
| Chronic pancreatitis, n [%] | 1 [7.1] | 1 [14.3] | 0 [0] |  |
| Any Complications within 30d, yes, n [%] | 12 [63.2] | 7 [100] | 5 [71.4] | 0.462 |
| Clavien–Dindo ≥ III, n [%] | 6 [42.9] | 4 [57.1] | 3 [42.9] | 0.225 |

### Metabolomic analysis

Gas chromatography–mass spectrometry was used to assess free fatty acids and monoacylglycerides in drain effluent samples, revealing pronounced compositional differences between CR-POPF and non-POPF cases. The heatmap of significantly altered fatty acids showed a clear segregation between CR-POPF and non-POPF drain effluent samples (Figure 1). Unsupervised principal component analysis (PCA) and supervised partial least squares discriminant analysis (PLS-DA), a class-based multivariate method maximizing separation between predefined groups, confirmed a robust separation between CR-POPF and control effluents, explaining more than 70 % of the total variance (Figure S1, S2). Variable-importance analysis identified palmitic acid, monopalmitin, and stearic acid as the top discriminating lipid species, indicating a consistent enrichment of long-chain saturated fatty acids in CR-POPF effluents (Figure S3). The volcano plot quantified these shifts, identifying multiple fatty acids with significant fold changes (FDR < 0.05; Figure 2). Monopalmitin was among the lipid species showing the most pronounced differences between CR-POPF and non-POPF samples (Figure S4). The quality-control evaluation demonstrated analytical stability, with missing values below 10% and uniform signal intensities across blank, control, and QC samples (Figure S5). Furthermore, correlation analysis revealed that palmitic acid levels correlated positively with other long-chain saturated fatty acids, including stearic, heptadecanoic, and myristic acid (Figure S6). This finding indicates a coordinated release of triglyceride-derived lipid species rather than isolated metabolite accumulation. Together, the metabolomic data defines a lipolytic lipid profile of CR-POPF effluents, enriched in long-chain saturated fatty acids and monoacylglycerides. Monopalmitin emerged as a robust marker of active lipolysis, showing consistent enrichment and high variable importance, and was therefore selected for downstream functional analyses.

**Figure 1.**
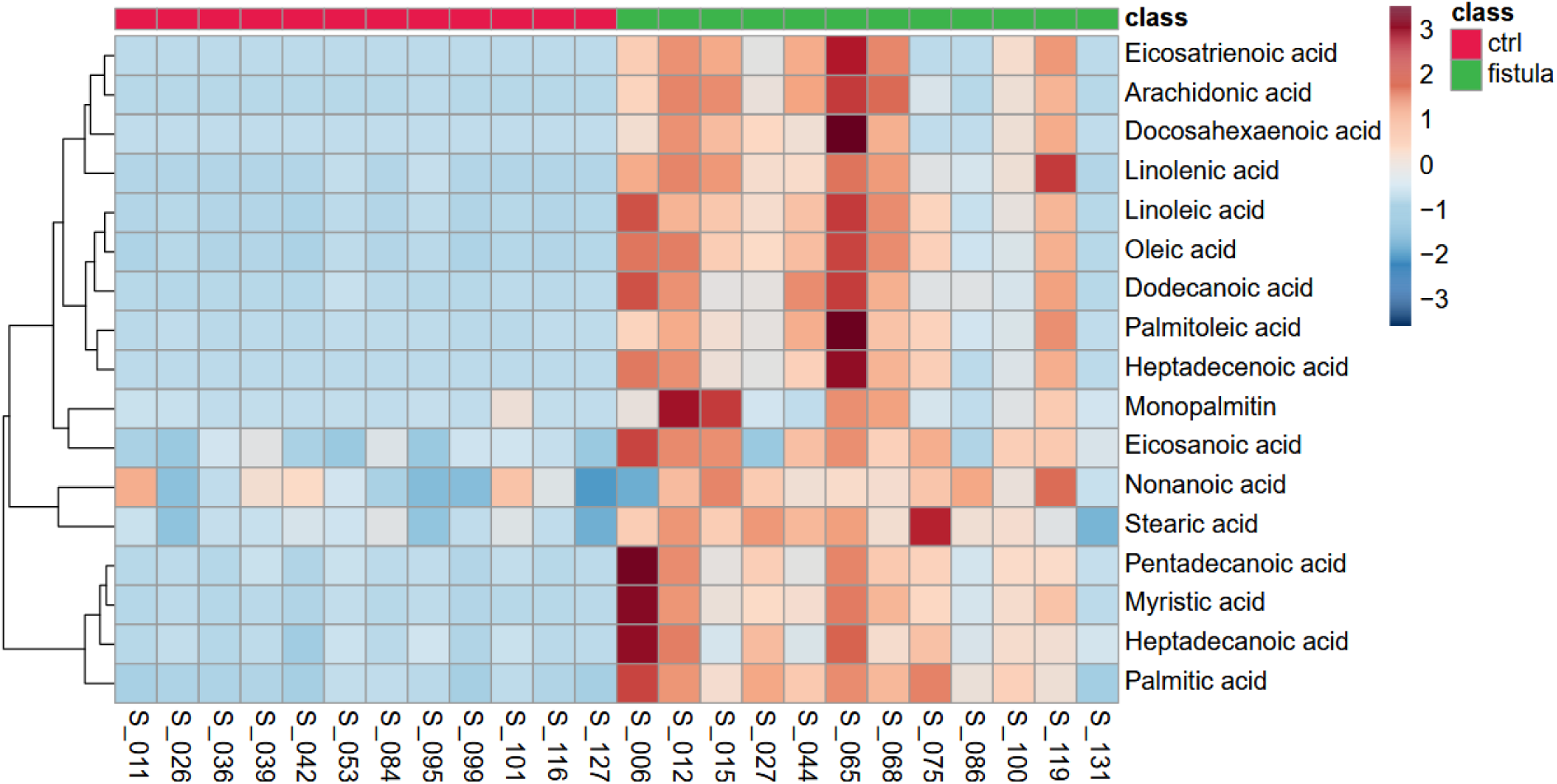
Heatmap of significantly altered fatty acids (FDR < 0.05). There is distinct clustering of CR-POPF versus control effluents with an enrichment of palmitic acid, monopalmitin, and stearic acid in the fistula group. Range-scaled z-scores are displayed.

**Figure 2:**
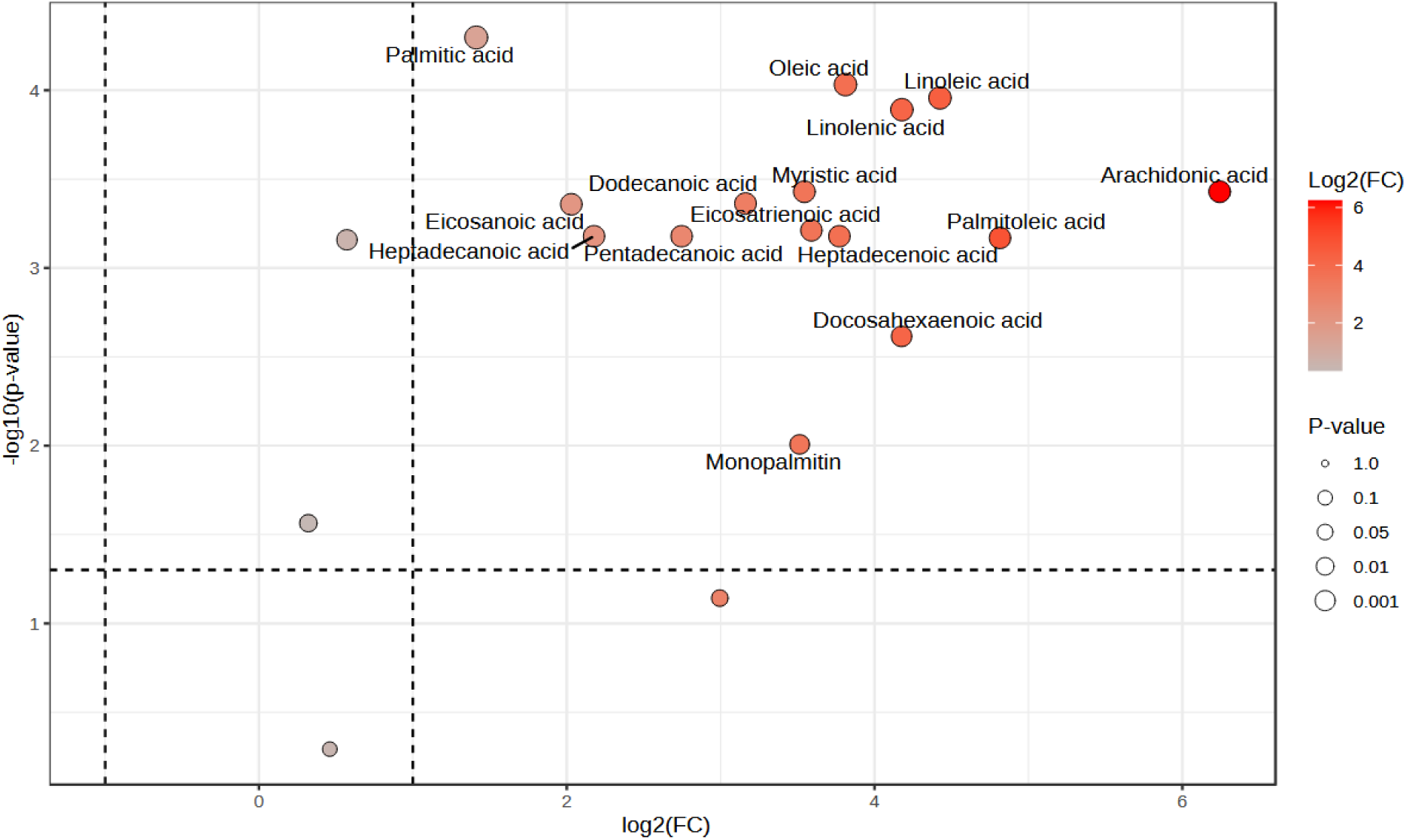
GC-MS metabolomic profiling identifies long-chain fatty acids that are enriched in CR-POPF effluents. Volcano plot summarizing differential abundance (log₂ fold change vs. −log₁₀ p value). Selected lipid species, including palmitic acid, monopalmitin, and stearic acid, are annotated in the volcano plot

### Fatty acid challenge across fibroblast, mesothelial, and carcinoma cell types

To establish subtoxic concentration ranges for subsequent experiments, HFF, peritoneal and pancreatic tumor cells were exposed to increasing concentrations of palmitic acid, stearic acid, and monopalmitin (0.1, 0.2, 0.5 mM). Two-way ANOVA revealed significant main effects of concentration and fatty acid type as well as significant interactions in all three cell types (HFF: F(8, 60) = 2.6, p = 0.0138; peritoneal: F(8, 57) = 12.8, p < 0.0001; PanC: F(8, 57) = 3.4, p = 0.0027). Repeated measures two-way ANOVA on normalized viability confirmed robust effects of both factors and their interaction (HFF: F(6, 36) = 44; peritoneal: F(6, 45) = 125; PanC: F(6, 45) = 34; all p < 0.0001). Across all models, concentration explained roughly one-third to one-half of the total variance, with the strongest contribution from fatty acid type. Viability decreased consistently with rising concentrations, most prominently at 0.5 mM. Palmitic acid caused the greatest loss of viability in all cell types, while stearic acid and monopalmitin produced only minor effects up to 0.2 mM. Collectively, these results defined 0.2 mM as the upper subtoxic concentration for the subsequent effluent-exposure and transcriptomic profiling experiments.

### Exposure to postoperative effluents: Pancreatic carcinoma cells (PanC-1)

In PanC-1 exposure to non-fistula effluents did not induce a relevant reduction in metabolic viability or an increase in membrane integrity–based cytotoxicity. Two-way ANOVA of absolute metabolic viability values showed significant effects of time (F(2,48) = 48.6, p < 0.0001) and treatment (F(7,48) = 7.1, p < 0.0001), but no interaction (F(14,48) = 0.15, p = 0.99). Analyses of normalized viability confirmed that variability was primarily driven by differences between individual non-fistula effluents (F(7,48) = 34.9, p < 0.0001), with no relevant time dependence. Overall, the ATP-based metabolic activity of PanC-1 pancreatic carcinoma cells, as assessed by CellTiter-Glo, remained close to that of the medium controls following exposure to non-fistula effluents. In contrast, exposure to CR-POPF effluents resulted in significant reductions in metabolic activity that varied between samples. Two-way ANOVA revealed significant main effects of time (F(2, 48) = 23.5, p < 0.0001) and treatment (F(7, 48) = 25.6, p < 0.0001), with no significant interaction effect (F(14, 48) = 0.48, p = 0.93). Treatment accounted for nearly two-thirds of the total variance and dominated the response pattern. Analyses of normalized metabolic activity confirmed that treatment was the only significant contributing factor (F(7, 48) = 229.7, p < 0.0001). Among the CR-POPF effluents, AES1448 and GR1479 were associated with the most significant reductions in metabolic activity at the analyzed time points and were therefore selected for further analysis (Figure S7).

### Exposure to postoperative effluents: Human fibroblasts (HFF)

Following exposure to non-fistula effluents, fibroblasts displayed stable ATP-based metabolic activity. A two-way ANOVA of the absolute metabolic activity values revealed significant main effects of time (F(2, 48) = 245.5, p < 0.0001) and treatment (F(7, 48) = 8.6, p < 0.0001), and no significant interaction effect (F(14, 48) = 0.82, p = 0.65). Analyses of normalized metabolic activity confirmed this pattern, with significant effects of time (F(2, 48) = 5.6, p = 0.0066) and treatment (F(7, 48) = 13.8, p < 0.0001). Metabolic activity remained close to medium controls across non-fistula effluents. In contrast, exposure to CR-POPF effluents resulted in pronounced, treatment-dependent reductions in metabolic activity. A two-way ANOVA revealed significant main effects of time (F(2, 48) = 34.0, p < 0.0001) and treatment (F(7, 48) = 32.5, p < 0.0001), but no significant interaction effect (F(14, 48) = 1.64, p = 0.10). Treatment explained approximately 62% of the total variance. In normalized analyses, treatment remained the only significant source of variation (F(7, 42) = 261.4, p < 0.0001). Among CR-POPF effluents, AES1448 and GR1479 were associated with the most pronounced and consistent reductions in metabolic activity across replicates and analyzed points of time (Figure S7).

### Exposure to postoperative effluents: Peritoneal mesothelial cells

Peritoneal cells exhibited stable ATP-based metabolic activity following exposure to non-fistula effluents. Two-way ANOVA of absolute metabolic activity values showed modest but significant main effects of time (F(2,48) = 8.1, p = 0.0009) and treatment (F(7,48) = 3.0, p = 0.0107), without a significant interaction (F(14,48) = 0.90, p = 0.998). Analyses of normalized metabolic activity confirmed these mild time-dependent changes (time F(2,48) = 3.8, p = 0.030; treatment F(7,48) = 5.1, p = 0.0002), with no effluent causing a pronounced reduction in metabolic activity relative to controls. In contrast, exposure to CR-POPF effluents resulted in a distinctly treatment-dependent response in peritoneal mesothelial cells. Two-way ANOVA demonstrated significant main effects of time (F(2,48) = 4.6, p = 0.015) and treatment (F(7,48) = 19.8, p < 0.0001), while the interaction remained non-significant (F(14,48) = 0.38, p = 0.97). Treatment explained nearly 70% of the total variance. Consistent with findings in fibroblasts and pancreatic carcinoma cells, the effluents AES1448 and GR1479 were associated with the most pronounced and reproducible reductions in metabolic activity across the analyzed time points, motivating their selection for subsequent transcriptomic analyses (Figure S7). Taken together, ATP-based metabolic activity assays indicated that non-fistula effluents were largely inert across all examined cell types, whereas only a subset of CR-POPF effluents induced consistent, treatment-dependent reductions in metabolic activity. Based on these findings, effluents AES1448 and GR1479, together with monopalmitin as a defined lipid reference stimulus, were selected for transcriptomic profiling in fibroblasts and peritoneal mesothelial cells.

### RNA sequencing and transcriptomic analysis

To delineate molecular programs engaged by lipolysis-derived fatty acids and CR-POPF effluents, mRNA sequencing was performed in non-malignant peritoneal mesothelial cell preparations and fibroblasts exposed to effluents AES1448, GR1479, monopalmitin (0.2 mM), or medium control. Transcriptomic profiling across all examined cell preparations revealed consistent yet stimulus-specific transcriptional responses to pancreatic effluents and lipolysis-derived fatty acids. Principal component analysis demonstrated a clear separation of experimental conditions along the dominant variance axis. Within individual peritoneal mesothelial cell preparations (P1–P6) and HFF, biological replicates formed compact clusters, indicating reproducible and stimulus-dependent transcriptional responses (Figure 3). Samples exposed to AES1448 and GR1479 clustered closer to medium controls, consistent with metabolically adaptive states, whereas monopalmitin exposure shifted the transcriptome toward a distinct expression space associated with proteotoxic and inflammatory stress responses. Differential expression analyses quantitatively confirmed the stimulus-dependent separations observed in PCA (Figure 4, S8, S9). Each stimulus induced a distinct transcriptional response pattern. To illustrate these stimulus-dependent patterns at the gene level, representative cell preparation–specific contrasts are shown. In peritoneal mesothelial cell preparation 5, exposure to AES1448 was associated with suppression of morphogenetic and developmental genes, including HOXA13, TBX5, and ALX4, alongside modest induction of metabolic and differentiation-associated transcripts such as ALDH1A3, WT1, BARX1, and HOXB7. GR1479 elicited a qualitatively similar but attenuated response, accompanied by moderate upregulation of metabolic genes including TRIM58 and SLPI. In contrast, monopalmitin exposure triggered a broad and relatively symmetric distribution of up- and downregulated genes, with prominent induction of stress- and lipid-handling transcripts such as ALDH1A3, HOXB4–7, HMOX1, CARD11, and SLPI. Across cell preparations, AES1448 and monopalmitin each modulated more than 6,000 transcripts, whereas GR1479 affected fewer, yet still substantial, gene sets. Functional annotation revealed two dominant transcriptional axes across conditions: a structural–transcriptional axis encompassing differentiation and cytoskeletal programs, and a metabolic–stress–associated axis reflecting oxidative stress responses and energy homeostasis. Together, these findings demonstrate that different stimuli elicit distinct modes of transcriptional activation. The observed differences suggest that variations in the underlying properties of the applied stimuli contribute to divergent cellular response patterns.

**Figure 3.**
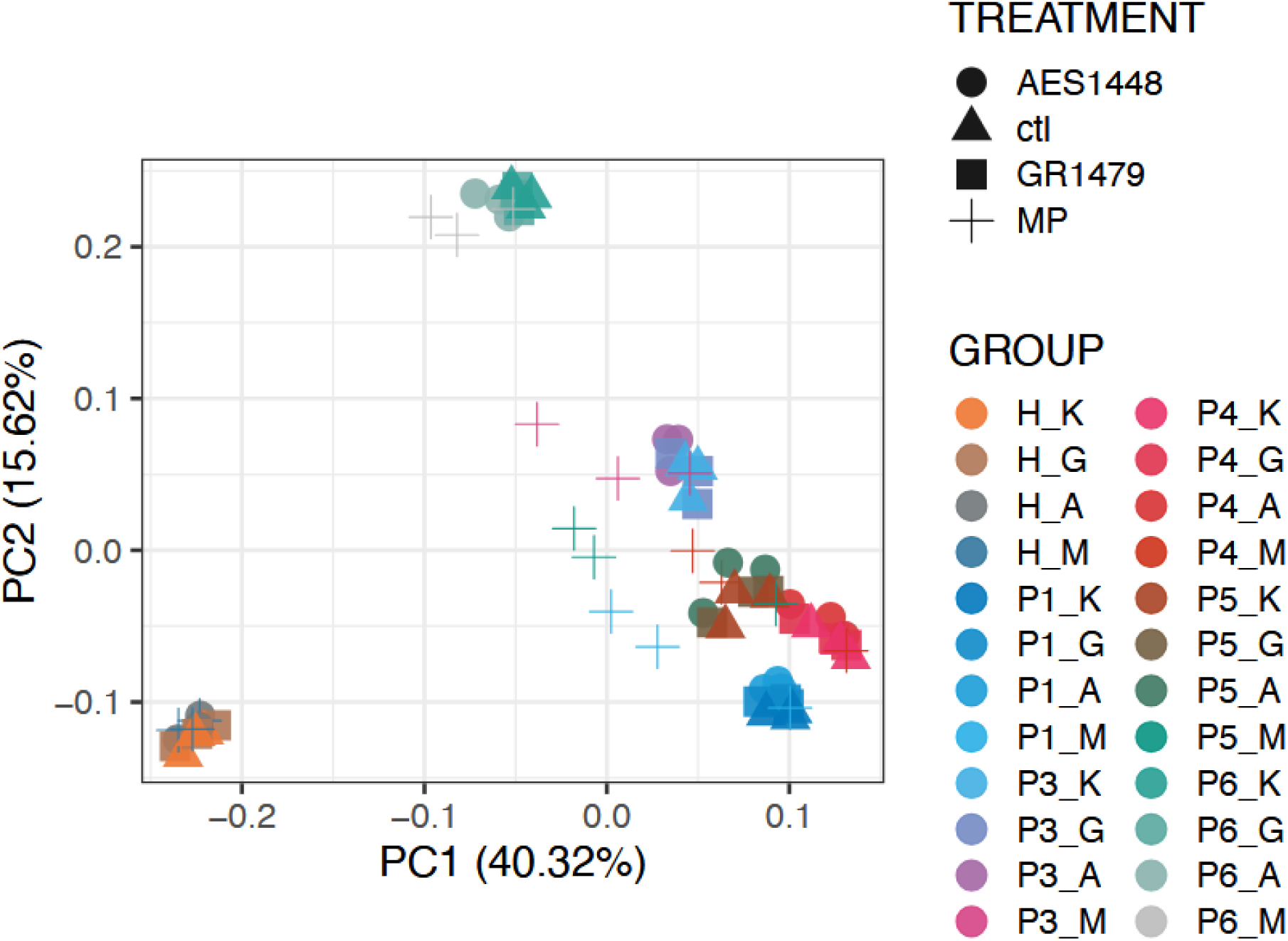
Principal component analysis of transcriptomic responses to lipolytic stimuli. PCA of TMM-normalized gene expression data. Points represent individual samples, colored by exposure condition (K, control; A, AES1448; G, GR1479; M, monopalmitin) and labeled by cell source. P1–P6 indicate primary peritoneal mesothelial cell preparations (one excluded after quality control), and H denotes the HFF fibroblast cell line. Replicates cluster tightly, indicating reproducible, stimulus-dependent transcriptional responses.

**Figure 4.**
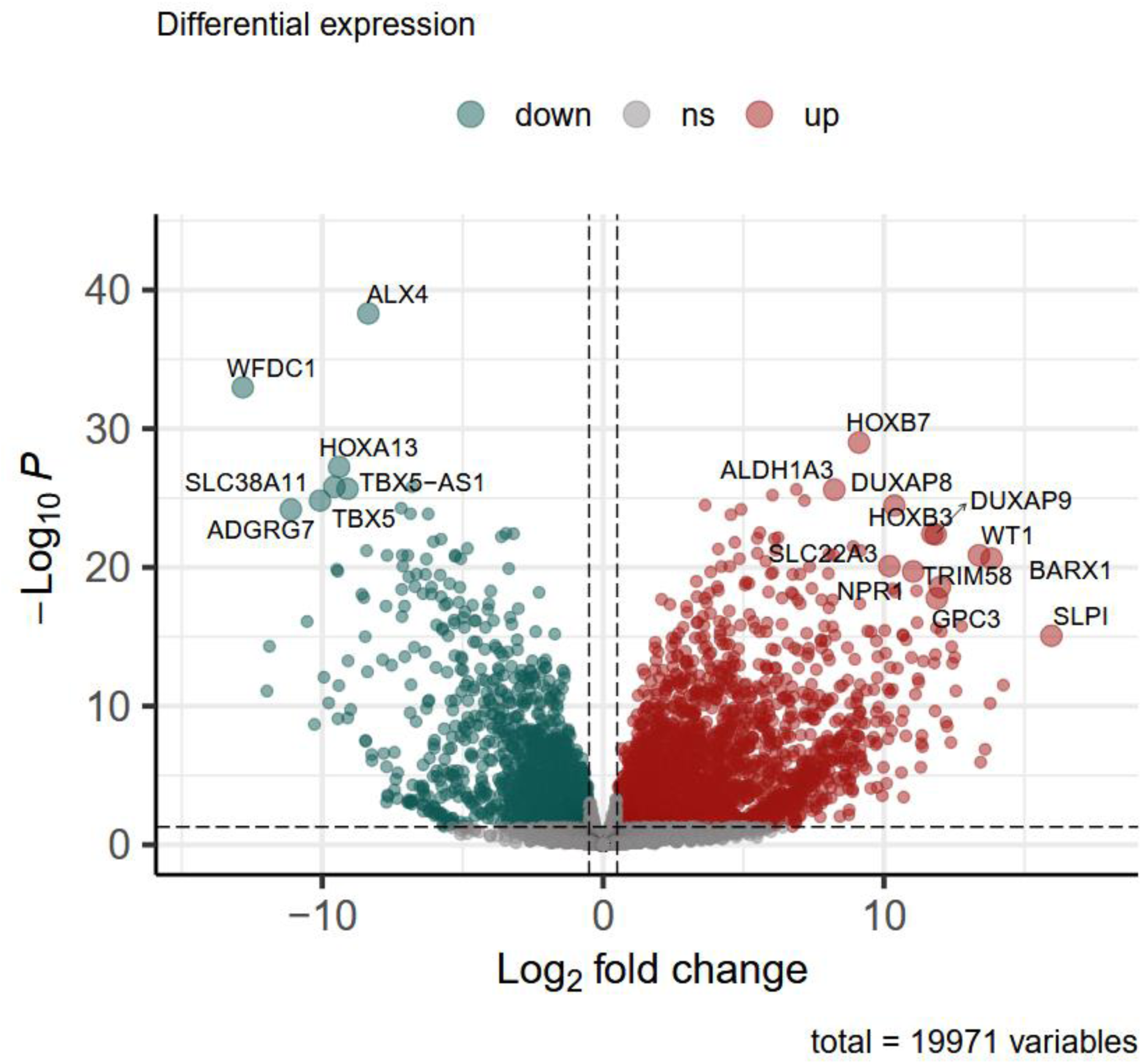
Cell source–specific differential gene expression under CR-POPF effluent exposure. Representative volcano plot comparing peritoneal mesothelial cell preparation 5 (P5) and human foreskin fibroblasts (HFF) following exposure to the CR-POPF effluent AES1448 under identical conditions. Genes are plotted by log₂ fold change (positive values indicate higher expression in P5; negative values indicate higher expression in HFF) and −log₁₀ adjusted p-value. Dashed lines denote the thresholds for differential expression (|log₂FC| ≥ 0.5, FDR < 0.05).

Gene set enrichment analyses translated these gene-level changes into functional pathway signatures. The number of significantly up- and down-regulated genes per treatment indicated more extensive transcriptional changes following exposure to AES1448 and monopalmitin compared with GR1479 (Figure S10). Hierarchical clustering of normalized enrichment scores from Hallmark GSEA divided the stimuli into two principal response groups: a metabolically adaptive phase (Control, GR1479) and a toxic-inflammatory phase (AES1448, Monopalmitin) (Figure 5). Under AES1448 exposure, *UV_RESPONSE_UP*and *INTERFERON_GAMMA_RESPONSE* were significantly enriched, while *Complement*, *Hypoxia*, *Glycolysis*, and *Interferon Alpha Response* trended upward. Conversely, *E2F Targets*, *DNA Repair*, *G2M Checkpoint*, and *Peroxisome* were repressed, indicating a combined stress–inflammation signature with reduced proliferative capacity. These transcriptional shifts comprised coordinated induction of DSG2, HOXB3/7, CARD11, and ALDH1A3, alongside suppression of E2F and RUNX family members, marking a transition from metabolic adaptation to oxidative–inflammatory imbalance.

**Figure 5.**
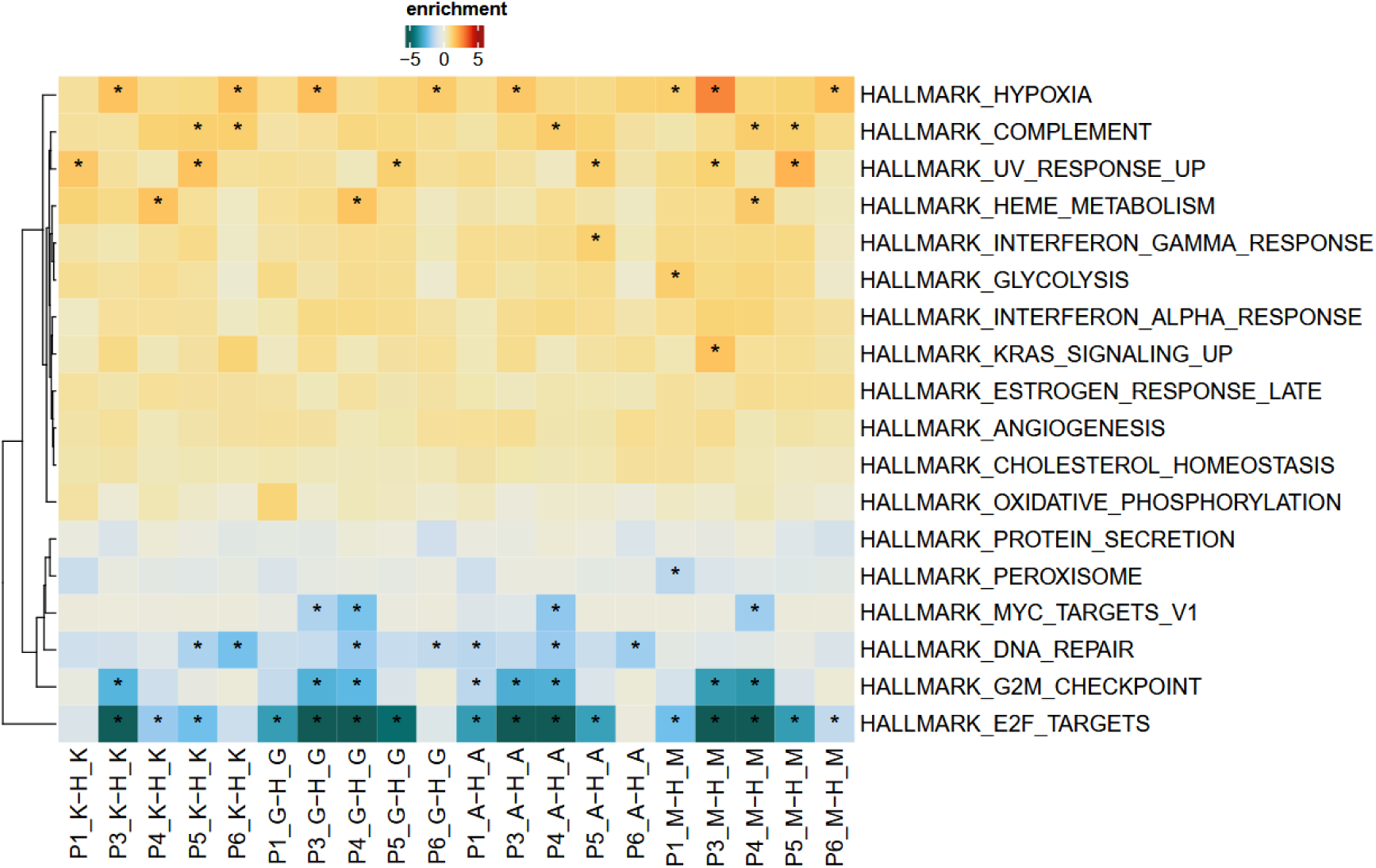
Hierarchical clustering of Hallmark GSEA results reveals distinct patterns of pathway enrichment across conditions. Control and GR1479 show comparatively attenuated and coherent enrichment profiles, whereas AES1448 and monopalmitin are associated with broader and more pronounced activation of stress- and metabolism-related Hallmark gene sets.

Condition-specific enrichment profiles further illustrate this progression (Figure 6). Exposure to AES1448 was associated with a combined immune and stress response, reflected by activation of pathways related to ultraviolet stress responses, interferon signaling, complement activation, and hypoxia, together with repression of cell-cycle–associated programs. This pattern is consistent with a pronounced inflammatory and stress-related transcriptional state. By contrast, GR1479 predominantly engaged pathways involved in mitochondrial energy metabolism and peroxisomal function, accompanied by only moderate attenuation of cell-cycle–associated programs (Figure S11). This transcriptional profile indicates a metabolically adaptive response without overt inflammatory activation.

**Figure 6:**
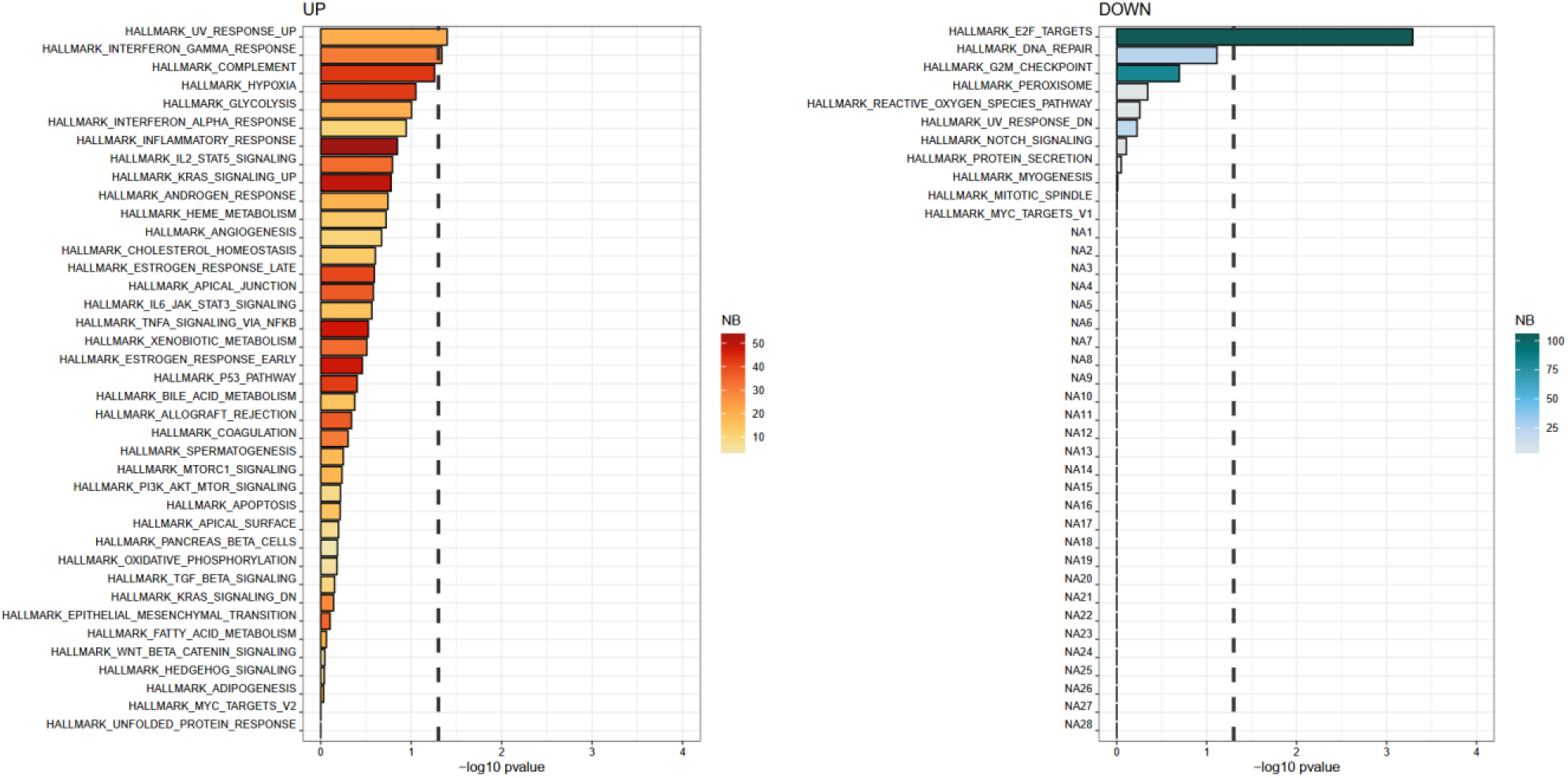
Condition-Specific Enrichment Profiles Reveal Different Transcriptional Responses to Pancreatic Effluents. AES1448 induces enrichment of the UV response up, interferon gamma response, complement, and hypoxia pathways, while repressing E2F targets and the G2M checkpoint. These results suggest a combined stress-inflammation response. The corresponding GSEA profiles of GR1479 and monopalmitin can be found in the Supplemental Material (Fig. S8A and S9).

Monopalmitin induced a distinct response marked by activation of hypoxia-, complement-, and stress-associated pathways, alongside suppression of cell-cycle and protein secretion programs. This constellation consists of advanced hypoxic-inflammatory stress and structural destabilization (Figure S12). These enrichment patterns delineate a secretion-dependent continuum of cellular states, ranging from metabolic adaptation (GR-like), through immune and stress activation (AES-like), to lipid-driven proteotoxic injury (monopalmitin).

To distinguish shared from stimulus-specific responses, pathway- and gene-level overlap analyses identified a compact core of shared stress-defense and condition-specific signaling modules (Figure 7A). Further intersection analyses clarified the extent of shared versus stimulus-specific transcriptional programs. These analyses identified approximately 140 up-regulated genes representing secretion-independent stress and defense pathways. These pathways are fundamental cytoprotective mechanisms that are likely triggered regardless of chemical composition (Figure 7B). Condition-specific enrichment patterns reflect distinct biological themes. Monopalmitin was dominated by endoplasmic reticulum (ER) stress and inflammatory signaling, AES1448 by immune differentiation and morphogenetic networks, and GR1479 by metabolic and peroxisomal reprogramming. At the gene level, the UpSet analysis illustrated quantitative overlaps between stimuli (Figure 7B). In peritoneal mesothelial cell preparation 5, monopalmitin induced the largest number of uniquely regulated genes (up = 638; down = 575), followed by AES1448 (up = 456; down = 462) and GR1479 (up = 433; down = 449). The greatest intersection occurred between AES1448 and GR1479 (up = 293), whereas monopalmitin partially overlapped with both (GR = 100; AES = 71). Extended pathway and GO analyses are provided in Supplemental Material (Figures S9-S13). These relationships remained consistent across cell preparations and replicates underscoring the robustness of the observed transcriptional architecture (Figure S17).

**Figure 7.**
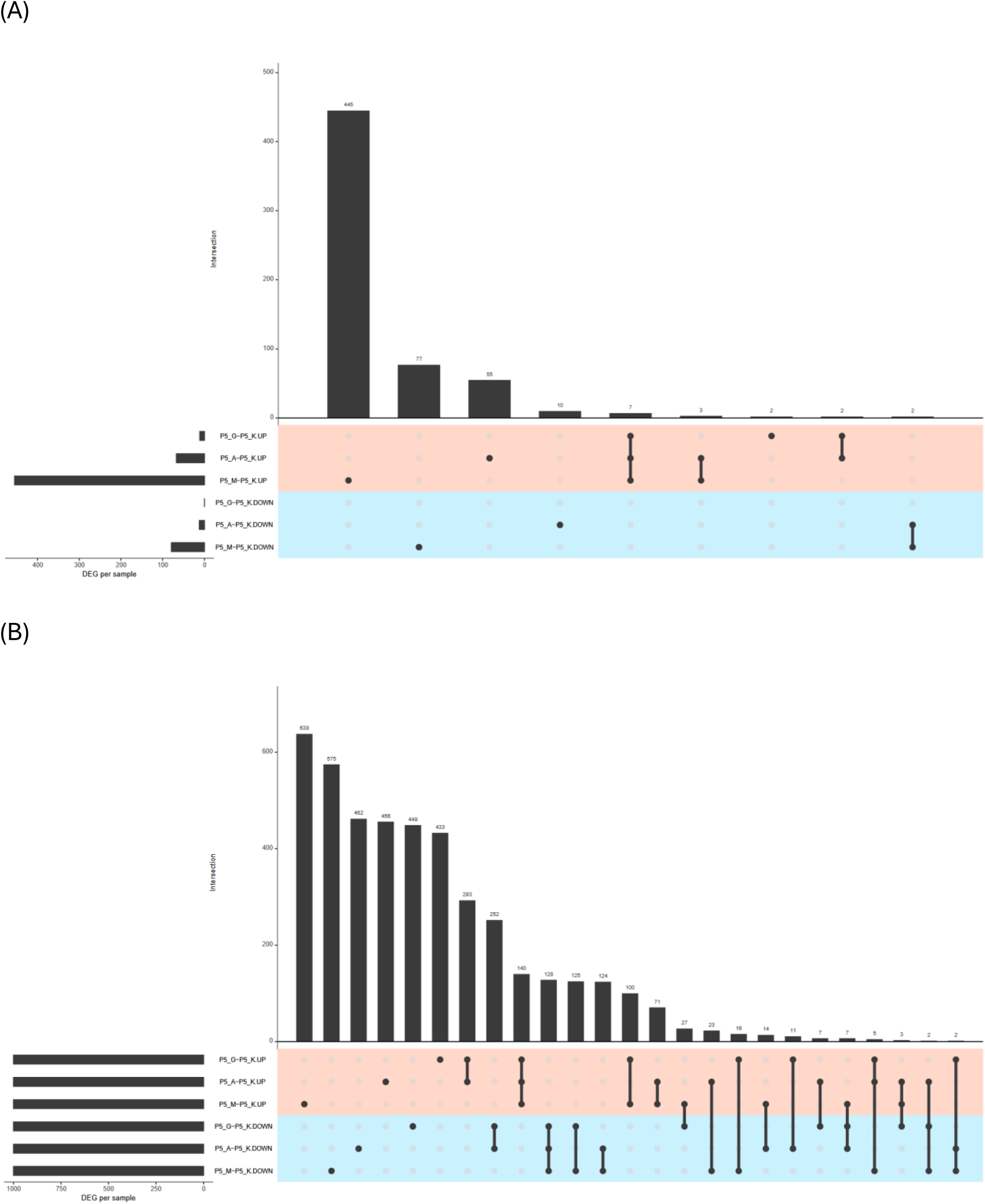
Shared and stimulus-specific transcriptional networks identified by intersection and enrichment analyses. Overlap analyses shown are representative of the observed patterns across preparations. **(A)** Fisher’s exact test of Hallmark pathway overlaps (top 10 terms), summarizing shared stress-defense and condition-specific signaling modules. **(B**) An UpSet plot showing the overlap between the top 1,000 differentially expressed genes in AES1448, GR1479, and Monopalmitin. There is strong overlap between AES1448 and GR1479, and the largest unique component is in Monopalmitin.

## Discussion

For years, the pathogenesis of CR-POPF was mainly interpreted in mechanical terms. However, growing evidence suggests a broader biological basis involving inflammation, premature enzyme activation, and lipotoxic effects (16,17).

Our findings align with this evolving concept. Distinct lipid compositions within postoperative pancreatic effluents determine their cytotoxic potential and provide evidence for patient-specific biochemical heterogeneity in CR-POPF pathogenesis. Using metabolomic and transcriptomic profiling, we identified fatty acid–driven injury patterns that link the composition of each effluent to stress responses in stromal and mesothelial cells. This integration shows that only certain effluents carry a lipotoxic signature that can trigger cell death and inflammation. This is consistent with the findings of Uchida et al., who identified intraperitoneal lipolysis as a key driver of postoperative inflammation and fistula progression. They also demonstrated that inhibiting lipase activity with drugs alleviates these effects (7).

To experimentally dissect these mechanisms, we established an in vitro model that replicates the stromal and mesothelial interface of the postoperative peritoneal cavity in accordance with current literature and standards (18). Our study design intentionally focused on this non-epithelial compartment rather than on tumor or acinar cells We excluded pancreatic ductal adenocarcinoma (PDAC) lines to avoid transcriptional bias from oncogenic signaling; such models do not reflect the postoperative tissue environment. Instead, we focused on peritoneal mesothelial cells and human fibroblasts, among the first non-malignant compartments exposed to pancreatic effluents after surgery and key cellular contributors to early anastomotic repair. This approach mirrors the early stromal–mesothelial interface implicated in peritoneal injury and healing, providing a physiologically relevant setting to study effluent-induced stress and inflammation independent of tumor-intrinsic pathways (19,20).

Monopalmitin was chosen as the reference stimulus among the lipolysis-derived candidates instead of free palmitic acid because of its higher solubility and physiological relevance. Monopalmitin is a monoglyceride generated during enzymatic triglyceride breakdown and reflects an amphiphilic lipid species present under lipolytic conditions. Owing to its high interfacial activity, it can modulate lipase binding at lipid–water interfaces and interact directly with cellular membranes (21). Leaking lipases from the pancreatic remnant can hydrolyze visceral fat, producing lipolytic products classically including free fatty acids and monoacylglycerides (22). Their accumulation in the intraperitoneal space provides a plausible biochemical link between uncontrolled lipolysis and local lipid toxicity, consistent with clinical and experimental observations (17).

This framework integrated three analytical layers: metabolomics, viability assays, and RNA sequencing. These layers were examined under tightly controlled lipid exposure. This sequential design directly links biochemical composition to functional toxicity and transcriptional response. This establishes a mechanistic bridge from effluent chemistry to cellular fate. Cytotoxicity did not represent a generic property of CR-POPF effluents. Instead, toxic effects arose from specific lipid components interacting with cell-type–specific vulnerability, thereby translating biochemical heterogeneity into distinct transcriptional and functional outcomes. Metabolomic profiling of drain effluents revealed a distinct lipid composition, dominated by long-chain saturated fatty acids (such as palmitic and stearic acid) and monoglycerides (such as monopalmitin). These lipid components accounted for most of the separation between CR-POPF and control effluents, clustering tightly within the fistula group. This profile is consistent with experimental models of pancreatitis, in which uncontrolled lipolysis of peripancreatic fat drives local and systemic inflammation, tissue necrosis, and maladaptive stromal remodeling, thereby shaping a disease-permissive microenvironment (9,23,24). Our data extends this principle to the postoperative setting. Our data suggests that local lipolytic activity alters the biochemical environment, which can lead to the development of clinically relevant pancreatic fistulas. Therefore, we examined how these biochemical signatures translate into cellular injury. To do so, we tested effluents and key lipid species across peritoneal mesothelial cells and fibroblasts, two central stromal compartments governing early anastomotic healing and repair. Viability assays in these cell models provided a biological context for the observed lipid patterns. Palmitic and stearic acids exhibited dose-dependent toxicity, establishing 0.2 mM as a subtoxic reference threshold. Notably, drain effluents were collected before the clinical manifestation of CR-POPF, allowing assessment of intrinsic biochemical toxicity independent of overt fistula severity. Within this range, most effluents were biologically inert. However, two CR-POPF samples (AES1448 and GR1479) consistently reduced viability across all tested cell types, indicating that cellular injury is driven by specific biochemical properties of individual effluents rather than by the presence of a fistula per se.

To link biochemical patterns to molecular programs, we performed RNA sequencing on a targeted subset of conditions: the two CR-POPF effluents that consistently reduced viability (AES1448 and GR1479) and the lipolysis-derived reference stimulus monopalmitin. This hypothesis-driven selection enabled high-resolution mapping of effluent-induced transcriptional stress programs. The first axis, observed with GR1479, reflects metabolic adaptation, including the upregulation of pathways related to oxidative phosphorylation, peroxisomal metabolism, and redox balance, which maintain partial cellular integrity. Consistent with previous findings, the second axis, which was prominent in AES1448 and monopalmitin, represented toxic inflammatory stress. This was characterized by an unfolded protein response, interferon signaling, and the suppression of proliferation and secretion modules (25,26). This dual-axis model illustrates the spectrum ranging from adaptive reprogramming to lipotoxic collapse. Several limitations should be acknowledged. First, the number of effluent samples was limited, reflecting the rarity of well-preserved CR-POPF collections. One sample was excluded from the viability analysis due to technical failure, which slightly reduced statistical power. In vivo validation is still pending, and deliberately excluding PDAC cell lines limits direct comparability to tumor-adjacent tissue. However, this approach ensures that the observed transcriptional programs reflect stromal and mesothelial responses that are more directly relevant to anastomotic healing, rather than tumor-intrinsic ones (27). Monopalmitin is a simplified yet physiologically plausible proxy for the complex CR-POPF lipid milieu.

Despite these limitations, the consistency across assays and cell types supports the robustness of the identified lipotoxic axis. Beyond its mechanistic relevance, this framework provides a biochemical foundation for translation by linking effluent composition to defined cellular transcriptional responses. In doing so, it delineates a potentially targetable lipotoxic pathway, opening avenues for pharmacological or perioperative strategies aimed at mitigating CR-POPF severity rather than merely predicting its occurrence. Growing evidence across pancreatology underscores the role of lipid metabolism and its toxic byproducts in pancreatic disease. Recent integrative approaches combining high-dimensional imaging, molecular profiling, and clinical data illustrate how AI-supported modeling can capture complex, non-linear disease mechanisms in related contexts (38,39). These examples highlight the broader applicability of multi-omics frameworks to pancreatic disorders characterized by fatty acid–driven injury (28,29).

Conceptually, CR-POPF should be viewed not solely as a mechanical failure of an anastomosis, but as a self-amplifying interaction between structural vulnerability, inflammation, and lipotoxic stress. Framing CR-POPF as a biochemical disease component superimposed on a structural lesion shifts the translational focus toward active mitigation of lipotoxic injury in the early postoperative phase. From this perspective, AI-based integrative modeling may serve as a tool to integrate drain lipid profiles, transcriptional stress signatures, and imaging features, thereby identifying high-risk biochemical constellations and informing mechanism-guided, locally targeted interventions. The successful implementation of such strategies will depend on harmonized biobanking, adequate quality assessment, FAIR-compliant data management, and prospective, multicenter validation.

## Data availability

The underlying metabolomic and transcriptomic datasets, as well as patient-level data not included in the Supplementary Materials, are available from the corresponding author upon reasonable request and subject to institutional and ethical regulations. RNA-seq data will be deposited in the Gene Expression Omnibus (GEO) and made publicly available prior to publication.

## Acknowledgment

We acknowledge financial support for J.D.L. from the Research Commission of the Faculty of Medicine, University of Freiburg. D.A.R. is supported by the German Research Foundation Deutsche Forschungsgemeinschaft, CRC1479 P17 - Project ID: 441891347 and by the German Cancer Aid, Deutsche Krebshilfe, Project ID: 70113697. G.A. is supported by EkoEstMed–FKZ 01ZZ2015.

## Author Contributions

J.D.L. contributed to conceptualization, methodology, data curation, writing of the original draft, review and editing, visualization, supervision, project administration, and funding acquisition. M.S. contributed to investigation, and manuscript review and editing. S.L. contributed to methodology, software, validation, formal analysis, investigation, resources, data curation, visualization, and manuscript review and editing. S.M. contributed to software, validation, investigation, supervision, and manuscript review and editing. S.C. contributed to data curation, manuscript review and editing, and project administration. S.F.-F. contributed to manuscript review and editing and project administration. G.A. contributed to methodology, software, validation, formal analysis, investigation, resources, data curation, visualization, and manuscript review and editing. B.K. contributed to methodology, software, validation, formal analysis, investigation, data curation, supervision, and manuscript review and editing. D.A.R. contributed to conceptualization, methodology, formal analysis, resources, manuscript review and editing, supervision, project administration, and funding acquisition. U.A.W. contributed to conceptualization, methodology, resources, manuscript review and editing, supervision, and project administration.

## Conflict of Interest Statement

The authors declare that the research was conducted in the absence of any commercial or financial relationships that could be construed as a potential conflict of interest.

## Author access and approval

All authors had full access to all data in the study and reviewed and approved the final version of the manuscript.

## Materials and Methods

### Data Collection and patient samples

Clinical data was obtained and entered in a prospective database. Patients who underwent pancreatic head resection for suspected PDAC between 2019 and 2025 were included. Postoperative abdominal drainage effluents were collected on the third postoperative day, prior to the clinical manifestation of CR-POPF. Patients were subsequently classified retrospectively into CR-POPF (n = 7) and non-POPF (n = 7) groups according to ISGPS criteria. The study was conducted in accordance with the Declaration of Helsinki and approved by the local institutional ethics committee (ID: 23-1302-S1). All procedures were conducted in accordance with the Declaration of Helsinki and Good Clinical Practice guidelines. The study adheres to the MDAR framework and its sequencing data comply with MINSEQE standards (30,31).

### Surgical Procedure of Pancreaticoduodenectomy

At our high-volume center, experienced pancreatic surgeons performed all pancreatic head resections as standard Whipple, pylorus-resecting or pylorus-preserving pancreaticoduodenectomy. In all cases analyzed, reconstruction consisted of duct-to-mucosa pancreatojejunostomy, hepaticojejunostomy and duodeno/ gastrojejunostomy. The jejunal loop was positioned in Child-S or Roux-en-Y orientation. One or two drains were routinely placed adjacent to the pancreatic anastomosis(32).

### Inclusion/Exclusion Criteria

Eligible patients were adults (≥18 years old) who underwent pancreatic head resection for suspected pancreatic ductal adenocarcinoma (PDAC) and had complete clinical, operative, and pathological data, as well as available postoperative drainage fluid within 72 hours after surgery. Patients were excluded if they underwent pancreatic resection for reasons other than PDAC, had a history of pancreatic surgery, had an intra-abdominal infection or sepsis at the time of resection, or had insufficient or contaminated drainage fluid samples (less than 2 mL, hemolysis, or bacterial overgrowth). Patients lost to follow-up within 30 days were also excluded.

### Patient Consent and Data Availability

All patients consented to use their anonymized data and samples for research purposes. Data were stored in a pseudonymized form according to institutional and national regulations. The datasets from this study are not public but available to the corresponding author upon reasonable request.

### Sex and gender reporting (SAGER guidelines)

This study was conducted in accordance with the Sex and Gender Equity in Research (SAGER) guidelines. Primary human peritoneal mesothelial cell preparations were derived from pseudonymized donors. No information on the biological sex or gender of these donors was available to the investigators at the time of experimentation, precluding sex-disaggregated analyses for these cell-based experiments. The same applies to the sex of the donors of the cell lines PanC-1 and human foreskin fibroblasts (HFF); no additional sex- or gender-specific information beyond vendor documentation was available or applicable for these in vitro models. For pancreatic effluent samples, donor sex was recorded and is reported in Table 1. Where relevant, sex-related information was therefore transparently disclosed for human-derived materials for which such data were available.

### Statistical analysis

Clinical and perioperative data were analyzed using SPSS (IBM SPSS Statistics, version 30; IBM Corp., Armonk, NY, USA) and in R (Version 4.4.0; R Core Team). Continuous variables are reported as median (interquartile range) or mean ± SEM, as appropriate. Group comparisons between patients with and without CR-POPF were performed using Student’s t test or Mann–Whitney U test for continuous variables and χ² test or Fisher’s exact test for categorical variables. A two-sided p value < 0.05 was considered statistically significant. For tables with more than two categorical levels, Pearson’s chi-square test (two-sided) was applied, whereas Fisher’s exact test was used for 2×2 tables. For in vitro assays, viability and cytotoxicity data were analyzed using two-way ANOVA with time and treatment as fixed factors. Where appropriate, repeated measures two-way ANOVA on normalized (% of control) values were used to account for within-experiment correlations. Post hoc comparisons were adjusted for multiple testing as indicated in the figure legends.

### Metabolomic Analysis

500 µl of pancreas drainage fluid was extracted by adding 500 µl chloroform containing 2 µg/ml methyl nonadecanoate as internal standard. After shaking for 5 min at 1400 rpm, samples were centrifuged for 10 min at 20,000 g and 4 °C. 200 µl of the lower phase were transferred into a new reaction tube and evaporated in a vacuum concentrator. Dried lipids were derivatized by adding 20 µl pyridine and 50 µl N-methyl-N-trimethylsilyl trifluoroacetamide and incubated for 30 min at 37°C and 1200 rpm. After 2 min centrifugation at 20,000 g at room temperature, 30 µl were transferred into a GC/MS vial and 30 µl of each sample were used to prepare a pooled quality control sample. Samples were injected in randomized order with regularly injected quality control samples in between using a GC-EI-MS equipped with an HP5-MS column as previously described (33). Intensities of each lipid were normalized to the internal standard and analyzed using the MetaboAnalyst 6.0 (34). Palmitic acid, monopalmitin, and stearic acid were selected for functional testing based on their enrichment profiles in the metabolomic screen. Monopalmitin arises from partial lipolysis of triglycerides and therefore reflects active lipase-mediated fat degradation rather than passive release of free fatty acids. Because of its higher aqueous stability and solubility, monopalmitin allowed reproducible dose–response testing in cell culture and was thus used as the primary stimulus for transcriptomic experiments.

### Cell lines and culture conditions

#### Human foreskin fibroblasts (HFF)

Human foreskin fibroblasts (HFF; ATCC® SCRC-1041™) were obtained from the American Type Culture Collection. Cells were cultured in Dulbecco’s Modified Eagle Medium (DMEM, high glucose, GlutaMAX) supplemented with 10% heat-inactivated fetal bovine serum (FBS, lot-matched), 1% penicillin–streptomycin, and 2 mmol/L L-glutamine at 37 °C in a humidified incubator with 5% CO₂. HFF were used for experiments at passages 6–10 and routinely tested negative for mycoplasma contamination.

#### Peritoneal mesothelial cells

Peritoneal mesothelial cells were isolated from omental tissue obtained from randomly selected non-oncological resection specimens preserved in the institutional biobank between 2020 and 2022. Only samples derived from the omentum majus were used for this study. Tissue collection and storage followed standardized biobanking protocols. Cells were isolated by enzymatic digestion and expanded in appropriate growth medium. Experiments were performed using five independent peritoneal mesothelial cell preparations at passages 3–7.

#### Pancreatic carcinoma cells (PanC-1 cell line)

Pancreatic carcinoma cells (PanC-1 cell line; ATCC® CRL-1469™), originally derived from a human pancreatic ductal adenocarcinoma (PDAC), were used as a tumor-derived epithelial cell line. Cells were maintained in DMEM and used at passages below 15. PanC-1 cells were excluded from transcriptomic analyses to avoid confounding effects of tumor-intrinsic genomic and transcriptional alterations and to focus on stromal cell–specific responses.

### Assays for cell viability and cytotoxicity

#### Luminescent cell viability assay (CellTiter-Glo®)

Cell viability was assessed using the CellTiter-Glo® Luminescent Cell Viability Assay (Promega, Madison, WI, USA), according to the manufacturer’s instructions. CellTiter-Glo quantifies intracellular ATP as a surrogate marker of metabolically active cells. Upon addition of the reagent containing luciferase and luciferin, cells are lysed and generate a stable luminescent signal (half-life > 5 h). Light intensity is directly proportional to the number of viable cells.

#### Fluorescence-based cytotoxicity assay (CellTox® Green)

Cytotoxicity was measured using the CellTox® Green Cytotoxicity Assay (Promega) according to the manufacturer’s instructions. CellTox Green monitors the loss of cell membrane integrity in real time. The assay employs an asymmetric cyanine dye that selectively penetrates compromised cell membranes and binds to free DNA, producing a fluorescent signal proportional to the number of dead cells.

#### Fatty acid challenge across stromal and epithelial cell types

HFF, PanC-1, and peritoneal mesothelial cell preparations were subjected to a series of dose–response experiments to assess cellular tolerance to individual fatty acids. Cells were cultured in serum-free medium supplemented with fatty acid–poor bovine serum albumin (BSA) as a carrier. Treatments included vehicle control (ethanol) and three lipid species, palmitic acid (PA), stearic acid (SA), and monopalmitin (MP), each complexed to BSA. Concentrations of 0.1 mM, 0.2 mM, and 0.5 mM were applied for 24 hours under standard culture conditions (37 °C, 5 % CO₂). After incubation, cell viability and cytotoxicity were assessed using standardized metabolic and membrane integrity–based assays. Data were summarized as mean ± SEM, and statistical testing was performed as indicated in the figure legends.

#### Exposure to postoperative effluents

In the second experimental step, cell viability and cytotoxicity assays were conducted using postoperative drain effluents obtained from patients with and without CR-POPF. Two experimental setups were established using non-fistula effluents and CR-POPF effluents, respectively. PanC-1, HFF, and peritoneal mesothelial cell preparations were exposed to patient-derived effluents diluted 1:5 (effluent: medium) in serum-free culture medium supplemented with fatty acid–poor bovine serum albumin (BSA). After equilibration, the culture medium was replaced with effluent-supplemented medium. Effluents from seven patients per group (non-fistula and CR-POPF) were used. For each condition, three biological replicates were performed per cell line. Cells were incubated with the respective effluents for 24, 48, or 72 hours. After each incubation period, cell viability and cytotoxicity were assessed using standardized metabolic and membrane integrity–based assays, and mean values per cell line were recorded.

#### RNA isolation and sequencing for transcriptomic profiling

Peritoneal mesothelial cell preparations were obtained from five independent non-malignant donors. Human foreskin fibroblasts (HFF) were used as a reference fibroblast cell line. Both cell types were exposed for 24 h to postoperative effluents selected based on prior viability screening. Effluents from two CR-POPF patients (AES1448 and GR1479) were used, alongside medium alone (CTRL) and monopalmitin (0.2 mM) as controls. For transcriptomic profiling, HFFs and peritoneal mesothelial cell preparations were re-exposed to the selected effluents diluted 1:50 (effluent: medium) in serum-free culture medium to ensure sufficient cell viability for RNA isolation. For each cell type (peritoneal cells and HFF) and condition (CTRL, AES1448, GR1479, monopalmitin 0.2 mM), three independent culture replicates were prepared. After exposure, total RNA was isolated using silica-membrane spin columns (RNeasy Mini Kit, Qiagen, Hilden, Germany) with on-column DNase digestion. RNA concentration and purity were determined spectrophotometrically, and integrity was verified by microcapillary electrophoresis; only samples with RIN ≥ 8 were used for sequencing. RNA sequencing libraries were prepared by Novogene GmbH (Planegg, Germany) using the TruSeq Stranded mRNA Library Prep Kit (Illumina) and sequenced on a NovaSeq 6000 platform (2 × 100 bp; ∼25–30 million reads per sample). Raw reads underwent quality control using FastQC and MultiQC, adapter trimming with Cutadapt, alignment to the human reference genome (GRCh38) using STAR, and gene-level read quantification with featureCounts. Differential expression analysis was performed on raw count data using DESeq2, applying a threshold of |log₂ fold change| ≥ 0.5 and a Benjamini–Hochberg–adjusted false discovery rate (FDR) < 0.05.

#### Transcriptomics analysis

Paired-end reads were trimmed to remove low-quality bases and adapter sequences using Trimmomatic (v0.39). edgeR was used for complementary differential expression and enrichment analyses to ensure robustness across normalization and statistical frameworks. Trimmed reads were aligned to the human reference genome (hg38), and reads per gene were quantified using STAR (v2.7.11a). Downstream analysis was performed in R (v4.4.0). Lowly expressed genes were removed using the filterByExpr function from the edgeR package, which retains only those genes with sufficient read counts (minimum of 10 counts in at least 70% of samples within a group, and a total count of at least 15 across all samples) to ensure reliable statistical inference. Counts were normalized by library size and the trimmed mean of M-values (TMM) method. Principal component analysis (PCA) of TMM-normalized log₂ counts per million (CPM) was performed to explore sample variance and clustering. Differential gene expression analysis was analyzed (i) between treatments within the same patient and (ii) across patients and healthy donor controls, with the Bioconductor package edgeR (v4.2.2). Genes were considered differentially expressed at |log₂ fold change| ≥ 0.5 and false discovery rate (FDR) < 0.05. Gene-set enrichment was performed using clusterProfiler (v4.12.6) with MSigDB as reference gene-sets. Enrichment scores were normalized (NES) and adjusted for multiple testing (FDR < 0.05). Heatmaps were generated to visualize expression patterns of top-ranked differentially expressed genes and enriched pathways. Keyword-based searches (“lipo”, “toxi”) were applied to enrichment outputs to highlight mechanistically relevant processes. To integrate results across conditions, Venn diagrams were constructed to illustrate overlaps between DEG sets, and significance of intersections was evaluated by Fisher’s exact test using the union of all differentially expressed genes as the reference universe. Supplementary materials include volcano plots, representative DEG lists, and detailed enrichment results corresponding to the main analyses.

## Abbreviations

AES1448: CR-POPF effluent sample (patient code)
ANOVA: Analysis of variance
BSA: Bovine serum albumin
CR-POPF: Clinically relevant postoperative pancreatic fistula
DEGs: Differentially expressed genes
DMEM: Dulbecco’s Modified Eagle Medium
FBS: Fetal bovine serum
FDR: False discovery rate
GC-MS: Gas chromatography–mass spectrometry
GSEA: Gene set enrichment analysis
GR1479: CR-POPF effluent sample (patient code)
HFF: Human foreskin fibroblasts
ISGPS: International Study Group on Pancreatic Surgery
MDAR: Materials, Design, Analysis and Reporting
MINSEQE: Minimum Information about a high-throughput Sequencing Experiment
MP: Monopalmitin
NES: Normalized enrichment score
NF-κB: Nuclear factor kappa-light-chain-enhancer of activated B cells
PA: Palmitic acid
PanC: Pancreatic carcinoma cells (Panc-1 line)
PCA: Principal component analysis
PDAC: Pancreatic ductal adenocarcinoma
POPF: Postoperative pancreatic fistula
QC: Quality control
RNA-seq: RNA sequencing
SA: Stearic acid
SPSS: Statistical Package for the Social Sciences
UPR: Unfolded protein response

## Notes

### Competing Interest Statement

The authors have declared no competing interest.

## References

1. Cameron JL, Riall TS, Coleman J, Belcher KA. One thousand consecutive pancreaticoduodenectomies. Ann Surg. 2006 Jul;244(1):10–5.

2. Theijse RT, Stoop TF, Hendriks TE, Suurmeijer JA, Smits FJ, Bonsing BA, et al. Nationwide Outcome after Pancreatoduodenectomy in Patients at very High Risk (ISGPS-D) for Postoperative Pancreatic Fistula. Ann Surg. 2023 Dec 11;281(2):322–8.

3. Bassi C, Marchegiani G, Dervenis C, Sarr M, Abu Hilal M, Adham M, et al. The 2016 update of the International Study Group (ISGPS) definition and grading of postoperative pancreatic fistula: 11 Years After. Surgery. 2017 Mar;161(3):584–91.

4. Khalid A, Amini N, Pasha SA, Demyan L, Newman E, King DA, et al. Impact of postoperative pancreatic fistula on outcomes in pancreatoduodenectomy: a comprehensive analysis of American College of Surgeons National Surgical Quality Improvement Program data. J Gastrointest Surg. 2024 Sep;28(9):1406–11.

5. Theijse RT, Stoop TF, Hendriks TE, Suurmeijer JA, Smits FJ, Bonsing BA, et al. Nationwide Outcome after Pancreatoduodenectomy in Patients at Very High Risk (ISGPS-D) for Postoperative Pancreatic Fistula. Annals of Surgery. 2025 Feb;281(2):322.

6. Eshmuminov D, Schneider MA, Tschuor C, Raptis DA, Kambakamba P, Muller X, et al. Systematic review and meta-analysis of postoperative pancreatic fistula rates using the updated 2016 International Study Group Pancreatic Fistula definition in patients undergoing pancreatic resection with soft and hard pancreatic texture. HPB (Oxford). 2018 Nov;20(11):992–1003.

7. Uchida Y, Masui T, Nakano K, Yogo A, Sato A, Nagai K, et al. Clinical and experimental studies of intraperitoneal lipolysis and the development of clinically relevant pancreatic fistula after pancreatic surgery. Br J Surg. 2019 Apr;106(5):616–25.

8. Nemecz M, Constantin A, Dumitrescu M, Alexandru N, Filippi A, Tanko G, et al. The Distinct Effects of Palmitic and Oleic Acid on Pancreatic Beta Cell Function: The Elucidation of Associated Mechanisms and Effector Molecules. Frontiers in Pharmacology. 2019;9.

9. Durgampudi C, Noel P, Patel K, Cline R, Trivedi RN, DeLany JP, et al. Acute Lipotoxicity Regulates Severity of Biliary Acute Pancreatitis without Affecting Its Initiation. Am J Pathol. 2014 Jun;184(6):1773–84.

10. Marchegiani G, Barreto SG, Bannone E, Sarr M, Vollmer CM, Connor S, et al. Postpancreatectomy Acute Pancreatitis (PPAP): Definition and Grading From the International Study Group for Pancreatic Surgery (ISGPS). Annals of Surgery. 2022 Apr;275(4):663.

11. Vlăduț C, Steiner C, Löhr M, Gökçe DT, Maisonneuve P, Hank T, et al. High prevalence of pancreatic steatosis in pancreatic cancer patients: A meta-analysis and systematic review. Pancreatology. 2025 Feb;25(1):98–107.

12. Patel K, Trivedi RN, Durgampudi C, Noel P, Cline RA, DeLany JP, et al. Lipolysis of Visceral Adipocyte Triglyceride by Pancreatic Lipases Converts Mild Acute Pancreatitis to Severe Pancreatitis Independent of Necrosis and Inflammation. Am J Pathol. 2015 Mar;185(3):808–19.

13. Liu Q, Gu X, Liu X, Gu Y, Zhang H, Yang J, et al. Long-chain fatty acids - The turning point between “mild” and “severe” acute pancreatitis. Heliyon. 2024 Jun 15;10(11):e31296.

14. Leishman S, Aljadeed NM, Qian L, Cockcroft S, Behmoaras J, Anand PK. Fatty acid synthesis promotes inflammasome activation through NLRP3 palmitoylation. Cell Rep. 2024 Aug 27;43(8):114516.

15. Xia W, Lu Z, Chen W, Zhou J, Zhao Y. Excess fatty acids induce pancreatic acinar cell pyroptosis through macrophage M1 polarization. BMC Gastroenterology. 2022 Feb 19;22(1):72.

16. Wüster C, Shi H, Kühlbrey CM, Biesel EA, Hopt UT, Fichtner-Feigl S, et al. Pancreatic Inflammation and Proenzyme Activation Are Associated With Clinically Relevant Postoperative Pancreatic Fistulas After Pancreas Resection. Ann Surg. 2020 Nov;272(5):863–70.

17. Nakamura N, Nagai K, Kaneda A, Yogo A, Kasai Y, Anazawa T, et al. Novel method to prevent severe postoperative pancreatic fistula caused by lipolysis. J Hepatobiliary Pancreat Sci. 2025 Jun;32(6):476–86.

18. Alsabeeh N, Chausse B, Kakimoto PA, Kowaltowski AJ, Shirihai O. Cell Culture Models of Fatty Acid Overload: Problems and Solutions. Biochimica et biophysica acta. 2017 Nov 15;1863(2):143.

19. Chen YT, Chang YT, Pan SY, Chou YH, Chang FC, Yeh PY, et al. Lineage Tracing Reveals Distinctive Fates for Mesothelial Cells and Submesothelial Fibroblasts during Peritoneal Injury. J Am Soc Nephrol. 2014 Dec;25(12):2847–58.

20. Mutsaers SE, Birnie K, Lansley S, Herrick SE, Lim CB, Prêle CM. Mesothelial cells in tissue repair and fibrosis. Front Pharmacol. 2015 Jun 9;6.

21. Reis P, Holmberg K, Miller R, Krägel J, Grigoriev DO, Leser ME, et al. Competition between Lipases and Monoglycerides at Interfaces. Langmuir. 2008 Jul 1;24(14):7400–7.

22. Stork E, Skeie S, Devold T, Østbø G, Flytkjær A, Jonsson E, et al. Enzymatic hydrolysis during *in vitro* digestion of triacylglycerols in a model system and milk: Isomerisation of diacylglycerols and monoacylglycerols. Food Chemistry Advances. 2025 Sep 1;8:101041.

23. Lilly AC, Astsaturov I, Golemis EA. Intrapancreatic fat, pancreatitis, and pancreatic cancer. Cell Mol Life Sci. 2023 Jul 15;80(8):206.

24. Okumura T, Ohuchida K, Sada M, Abe T, Endo S, Koikawa K, et al. Extra-pancreatic invasion induces lipolytic and fibrotic changes in the adipose microenvironment, with released fatty acids enhancing the invasiveness of pancreatic cancer cells. Oncotarget. 2017 Feb 17;8(11):18280–95.

25. Moncan M, Mnich K, Blomme A, Almanza A, Samali A, Gorman AM. Regulation of lipid metabolism by the unfolded protein response. J Cell Mol Med. 2021 Feb;25(3):1359–70.

26. Volmer R, van der Ploeg K, Ron D. Membrane lipid saturation activates endoplasmic reticulum unfolded protein response transducers through their transmembrane domains. Proc Natl Acad Sci U S A. 2013 Mar 19;110(12):4628–33.

27. Lin YC, Hou YC, Wang HC, Shan YS. New insights into the role of adipocytes in pancreatic cancer progression: paving the way towards novel therapeutic targets. Theranostics. 2023 Jul 3;13(12):3925–42.

28. Cheng S, Hu G, Zhang S, Lv R, Sun L, Zhang Z, et al. Machine Learning-Based Radiomics in Malignancy Prediction of Pancreatic Cystic Lesions: Evidence from Cyst Fluid Multi-Omics. Adv Sci (Weinh). 2025 Apr 28;12(20):2409488.

29. Wang G, Yao H, Gong Y, Lu Z, Pang R, Li Y, et al. Metabolic detection and systems analyses of pancreatic ductal adenocarcinoma through machine learning, lipidomics, and multi-omics. Science Advances. 2021 Dec 22;7(52):eabh2724.

30. Brazma A, Hingamp P, Quackenbush J, Sherlock G, Spellman P, Stoeckert C, et al. Minimum information about a microarray experiment (MIAME)-toward standards for microarray data. Nat Genet. 2001 Dec;29(4):365–71.

31. Chambers K, Collings A, Graf C, Kiermer V, Mellor DT, Macleod MR, et al. Towards minimum reporting standards for life scientists [Internet]. MetaArXiv; 2019 [cited 2026 Feb 12]. Available from: https://osf.io/9sm4x

32. Warren KW, Cattell RB. Basic Techniques in Pancreatic Surgery. Surgical Clinics of North America. 1956 Jun 1;36(3):707–24.

33. Lagies S, Pichler R, Kaminski MM, Schlimpert M, Walz G, Lienkamp SS, et al. Metabolic characterization of directly reprogrammed renal tubular epithelial cells (iRECs). Sci Rep. 2018 Mar 1;8(1):3878.

34. Pang Z, Lu Y, Zhou G, Hui F, Xu L, Viau C, et al. MetaboAnalyst 6.0: towards a unified platform for metabolomics data processing, analysis and interpretation. Nucleic Acids Res. 2024 Jul 5;52(W1):W398–406.

