## Supplemental Material for "Lipotoxic fingerprints in clinically relevant postoperative pancreatic fistula: fatty acid–driven cytotoxicity targets cells involved in anastomotic healing"

**An in vitro experimental study.**

### Supplemental Material

1 Department of General and Visceral Surgery, Center for Surgery, Medical Center University of Freiburg, Hugstetterstraße 55, 79106 Freiburg, Germany

2 German Cancer Consortium (DKTK), Partner Site Freiburg and German Cancer Research Center (DKFZ), Heidelberg, Germany.

3 BIOSS Center of Biological Signaling Studies, University of Freiburg, Schänzlestaße 18, 79104 Freiburg, Germany

4 Core Competence Metabolomics, Hilde-Mangold-Haus, University of Freiburg, 79104 Freiburg, Germany

5 Spemann Graduate School of Biology and Medicine (SGBM), University of Freiburg, 79104 Freiburg, Germany

6 Institute of Medical Bioinformatics and Systems Medicine, Medical Center University of Freiburg, Faculty of Medicine, University of Freiburg, Freiburg, Germany

† Equal Contribution, shared last Authorship

### MDAR Reporting Checklist

| Domain | Item | Response |
|---|---|---|
| **Materials** | Newly created materials | No new materials generated; all analytes (lipids) commercially available. Not applicable. |
| | Antibodies | Not applicable. No antibodies used. |
| | DNA and RNA sequences | Not applicable. No novel sequences generated; sequencing performed on endogenous transcripts only. |
| | Cell materials | Human fibroblast (HFF-1), peritoneal mesothelial cells, and pancreatic ductal epithelial cells (HPDE) from certified repositories (ATCC). Mycoplasma-negative. |
| | Experimental animals | Not applicable. No in-vivo work performed. |
| | Plants and microbes | Not applicable. |
| | Human research participants | Drain effluents obtained from 14 surgical patients after informed consent; demographics and ethics approval reported in Methods. |
| **Design** | Study protocol | Mechanistic multi-omics study; not preregistered (exploratory translational design). |
| | Laboratory protocol | Full protocols for GC-MS and RNA-seq provided in Methods; standard references cited. |
| | Sample size determination | Not formally powered; exploratory study with n = 14 patient samples. |
| | Randomization | Cell treatments randomized across plates. |
| | Blinding | Investigators blinded to effluent group during viability assays and RNA-seq library preparation. |
| | Inclusion/exclusion criteria | One effluent was excluded due to technical measurement failure; PDAC cell lines were deliberately excluded to avoid oncogenic bias. |
| | Sample definition and replication | Viability and cytotoxicity assays were performed in triplicate (biological n = 3 per condition). RNA-seq analyses included three biological replicates per group (RIN ≥ 8; 25–30 M reads/sample). |
| | Ethics | Approved by institutional ethics committee; informed consent |

| | | |
|---|---|---|
| | | obtained from all participants (Ethics ID: 23-1302-S1). |
| | Dual Use Research of Concern (DURC) | Not applicable. |
| **Analysis** | Attrition | One effluent excluded due to technical measurement failure (2-way ANOVA dataset). |
| | Statistics | Two-way ANOVA (stimulus × concentration) for viability/cytotoxicity; edgeR for differential expression ($FDR < 0.05$); GSEA for pathway enrichment. Detailed workup included in Methods. |
| | Data normalization / QC | GC-MS data were normalized to internal standards and total ion count. RNA-seq data were processed as described in the RNA-seq statistical analysis section. |
| | Data availability | RNA-seq data were processed and curated in accordance with ENCODE and MDAR standards. GC–MS data were generated and curated following the Metabolomics Standards Initiative (MSI) guidelines. RNA-seq datasets will be deposited in the Gene Expression Omnibus (GEO) and made publicly available upon publication. Additional processed data supporting the findings of this study are available from the corresponding author upon reasonable request and in accordance with institutional and ethical regulations. |
| | Code availability | Custom R scripts supporting the analyses (R v4.4.0; edgeR, clusterProfiler) are available from the corresponding author upon reasonable request. |
| **Reporting** | Adherence to community standards | Study follows MDAR for transparency. MDAR checklist provided in Supplementary Information. |

### Metabolomic Analysis

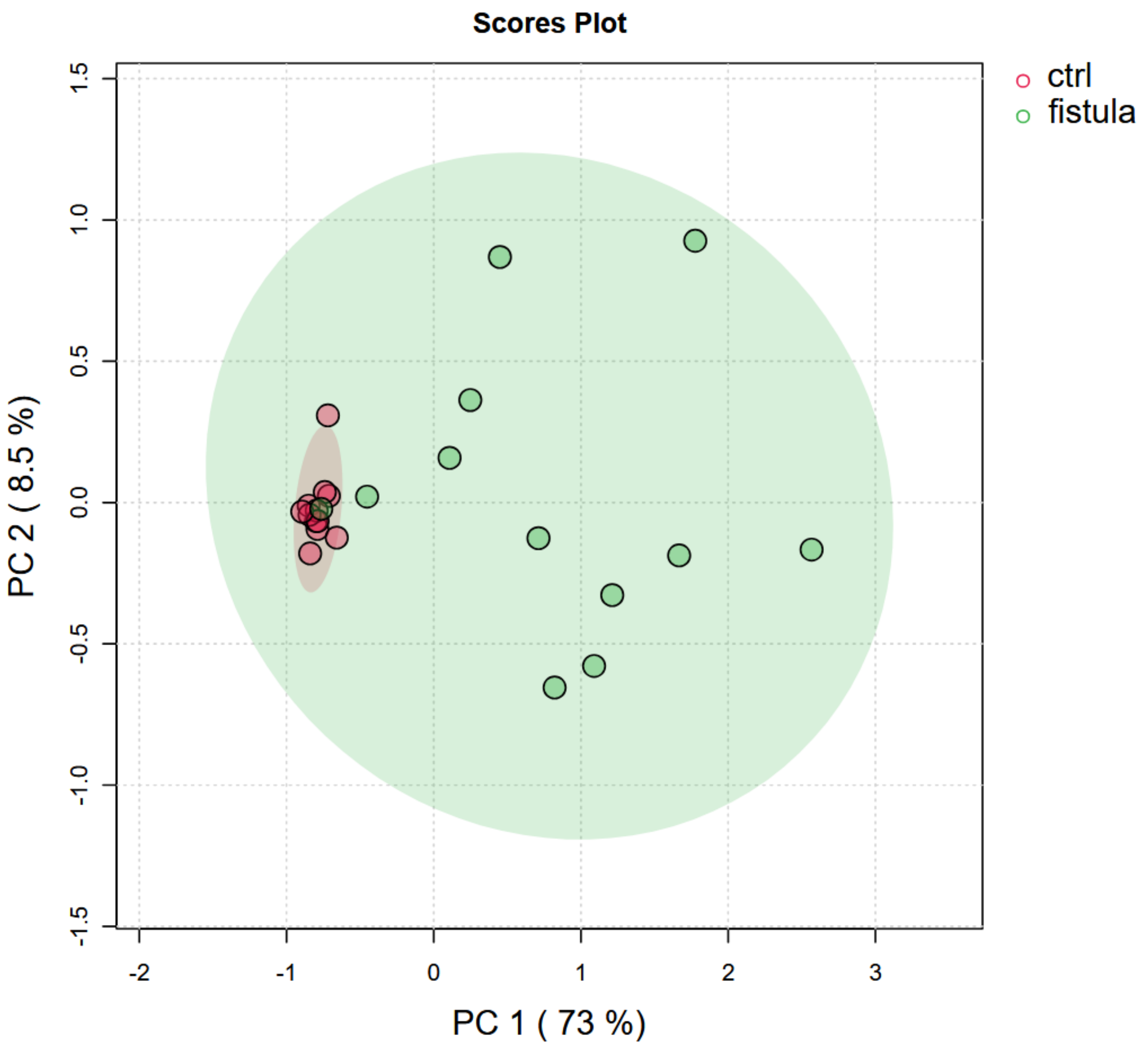


***Figure S1. Principal component analysis (PCA) of control and CR-POPF samples.*** *Unsupervised PCA scores plot showing the distribution of control and CR-POPF samples along the first two principal components (PC1 and PC2). Shaded ellipses represent 95% confidence regions for each group.*

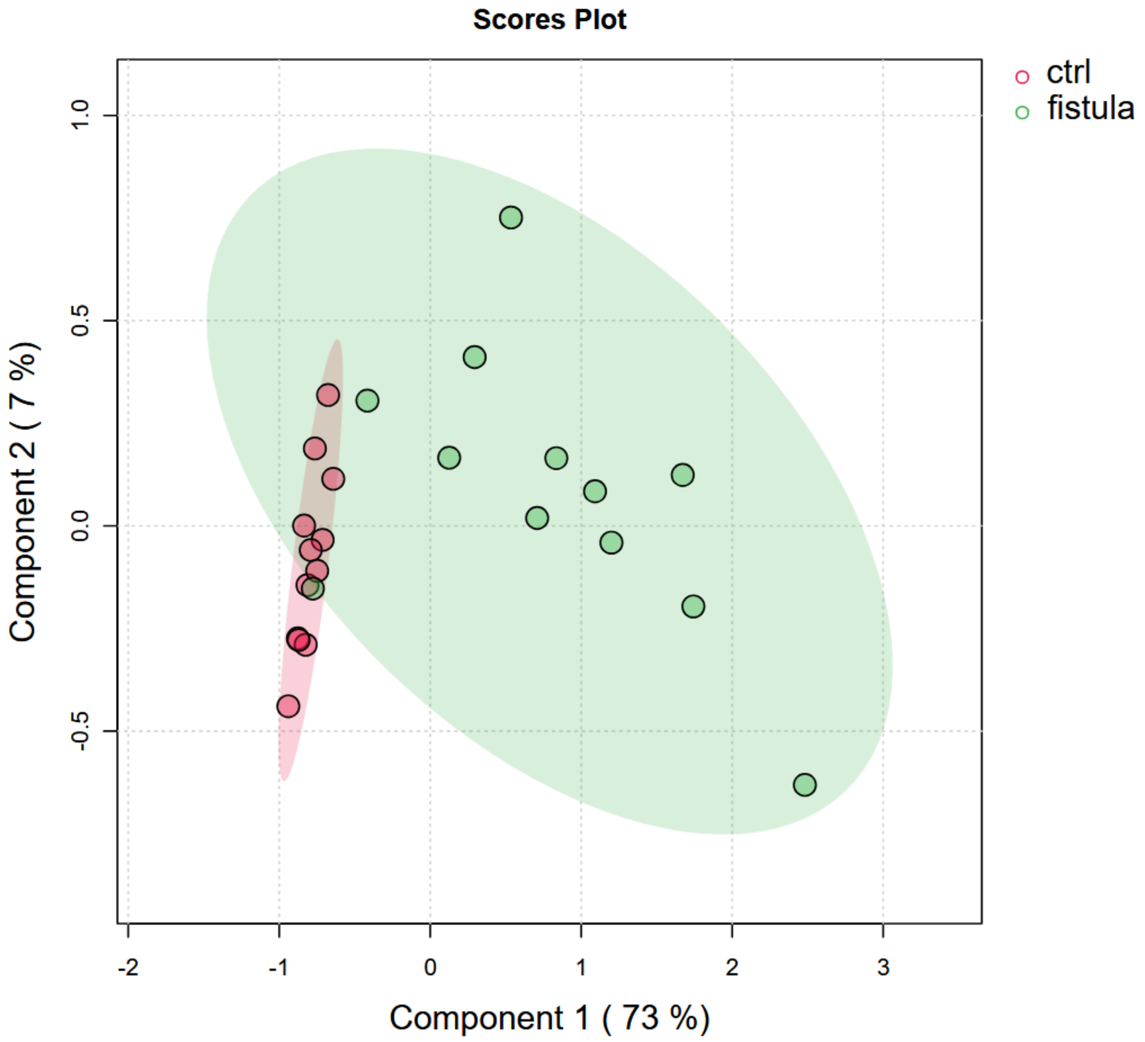


***Figure S2. Partial least squares discriminant analysis (PLS-DA) of control and CR-POPF samples.*** *PLS-DA scores plot showing the distribution of control and CR-POPF samples along the first two latent components. Shaded ellipses represent 95% confidence regions for each group.*

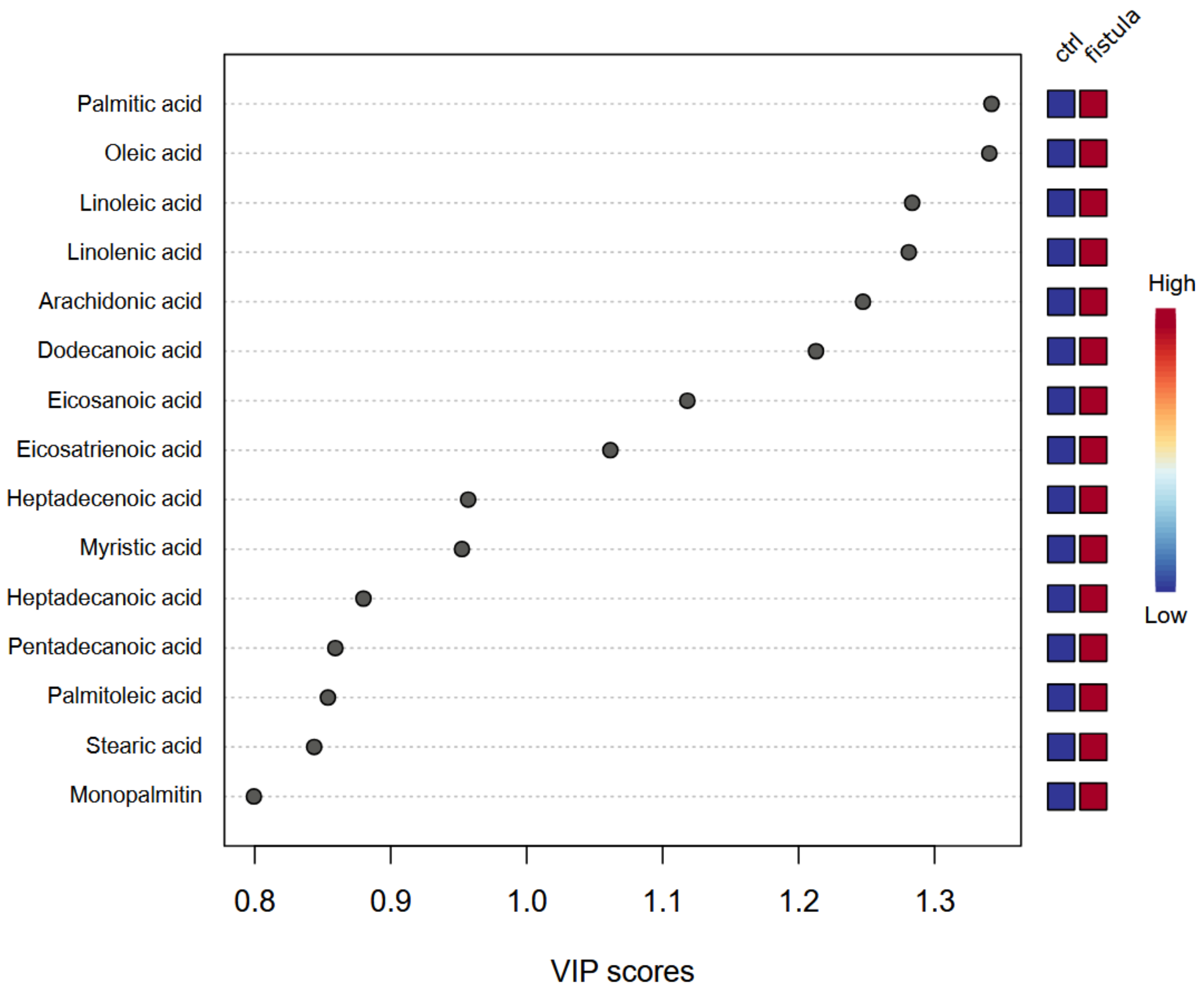


***Figure S3. Variable importance in projection analysis.*** *Lipid species are ranked by VIP scores based on their contribution to discrimination between control and fistula samples. Long-chain fatty acids showed the highest importance for group separation, whereas monoglycerides contributed less to discrimination despite their functional relevance in downstream analyses.*

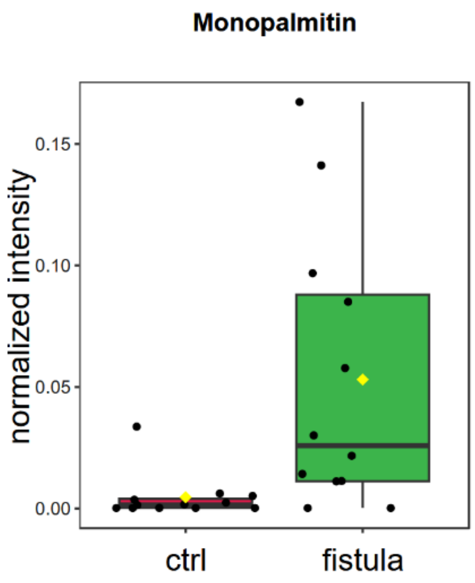


***Figure S4. Box–swarm plot of monopalmitin levels.*** *Normalized monopalmitin intensities are shown for control and CR-POPF drain effluent samples. Boxes represent the median and interquartile range, with individual measurements overlaid. The between-group difference reached nominal statistical significance (unadjusted p < 0.05).*

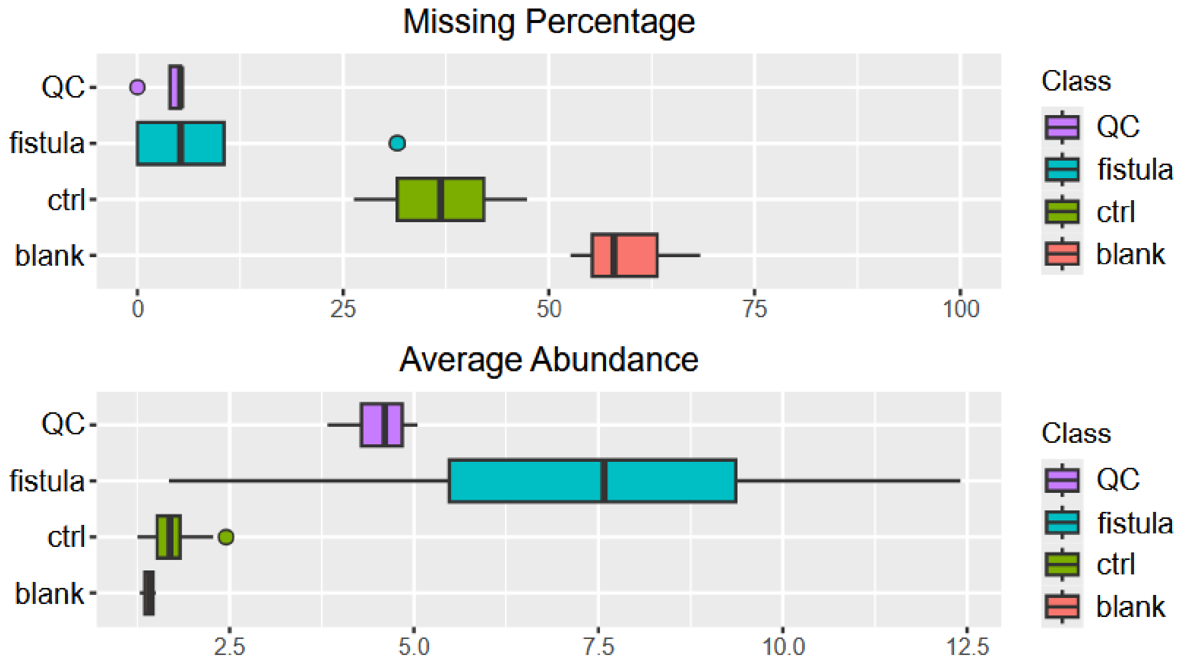


***Figure S5. Summary of the quality control metrics for the GC-MS analysis of postoperative drain effluents.*** *This figure shows the percentage of missing values across blank, control, CR-POPF, and pooled QC samples. The average signal abundance per class demonstrates a uniform instrument response. The stable intensity distributions of regularly injected pooled QC samples throughout the run confirm analytical reproducibility.*

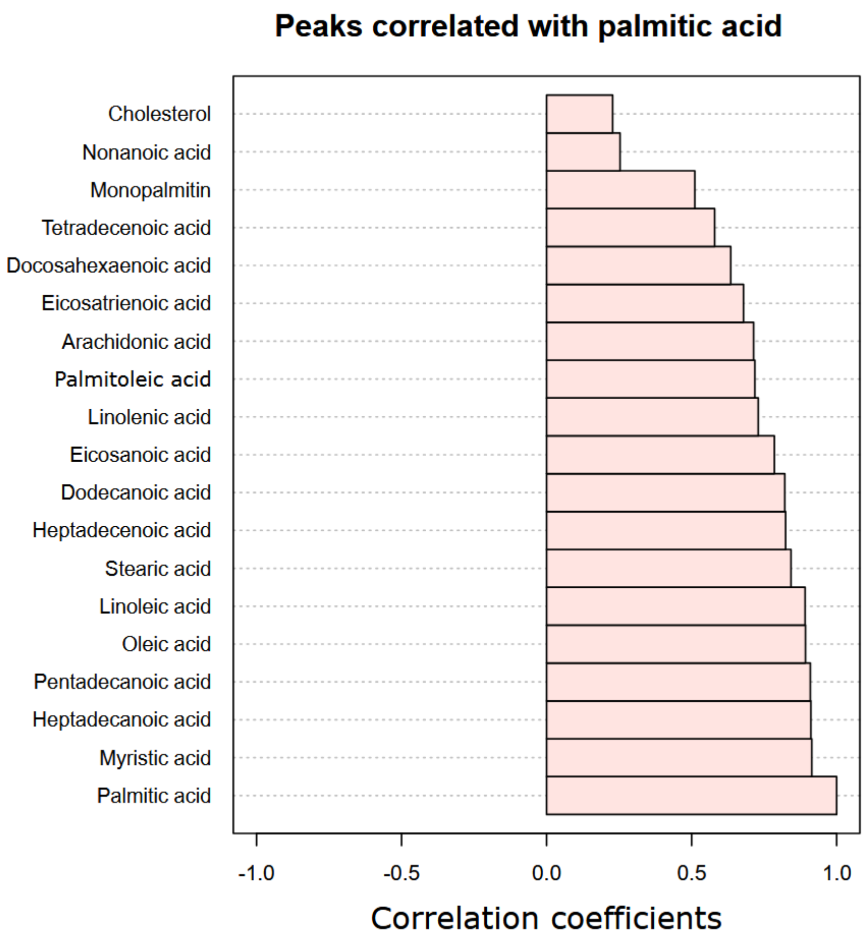


***Figure S6. Correlation Analysis of Quantified Fatty Acids.*** *This figure illustrates the co-enrichment of long-chain saturated fatty acids (e.g., stearic acid, monopalmitin, and heptadecanoic acid) with palmitic acid using Pearson correlation coefficients.*

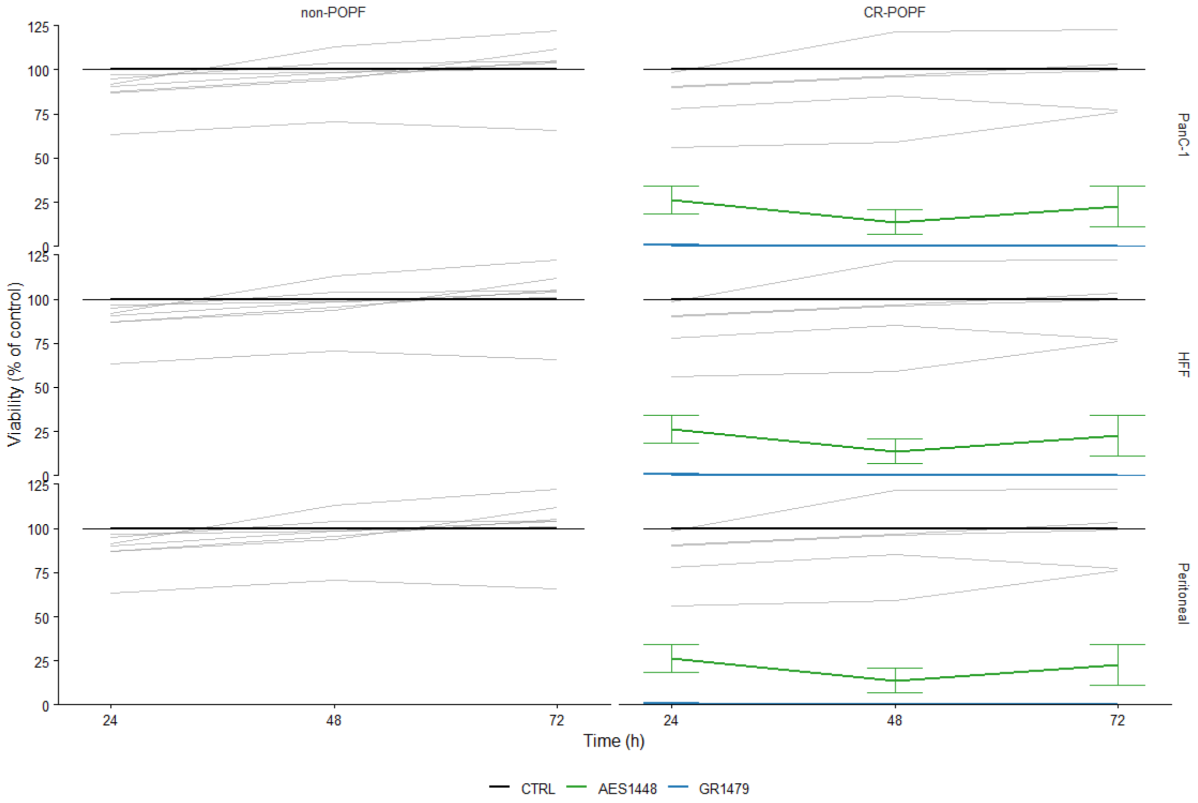


***Figure S7. Postoperative effluents and cellular metabolic viability.*** *Pancreatic carcinoma cells (PanC-1), human foreskin fibroblasts (HFF), and peritoneal mesothelial cells were exposed to non-POPF or CR-POPF effluents for 24, 48, and 72 h. The CR-POPF effluents AES1448 and GR1479, which showed the most divergent metabolic profiles, are shown separately. Cellular metabolic viability was quantified using an ATP-based assay and is expressed relative to control medium (CTRL). Data are presented as mean ± SEM from three independent experiments.*

**Transcriptomic Analysis**

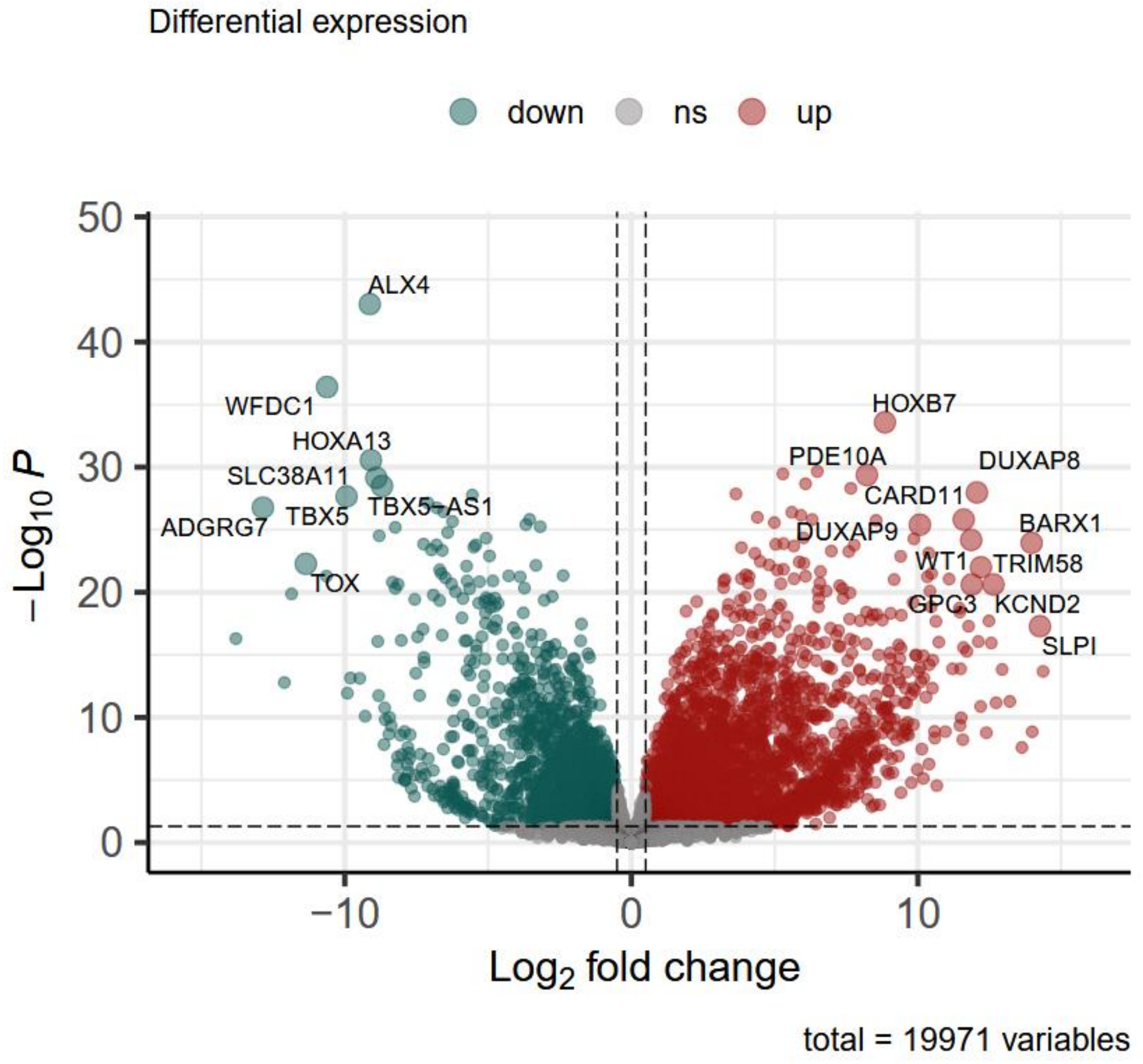


***Figure S8. Stimulus-specific differential gene expression under GR1479 exposure.*** *Representative volcano plot comparing peritoneal mesothelial cell preparation 5 (P5) following exposure to GR1479 with matched control conditions. Genes are plotted by $\log_2$ fold change (positive values indicate higher expression in GR1479-exposed cells; negative values indicate higher expression in controls) and $-\log_{10}$ adjusted p-value. Dashed lines denote the thresholds for differential expression ($|\log_2 FC| \geq 0.5$, $FDR < 0.05$).*

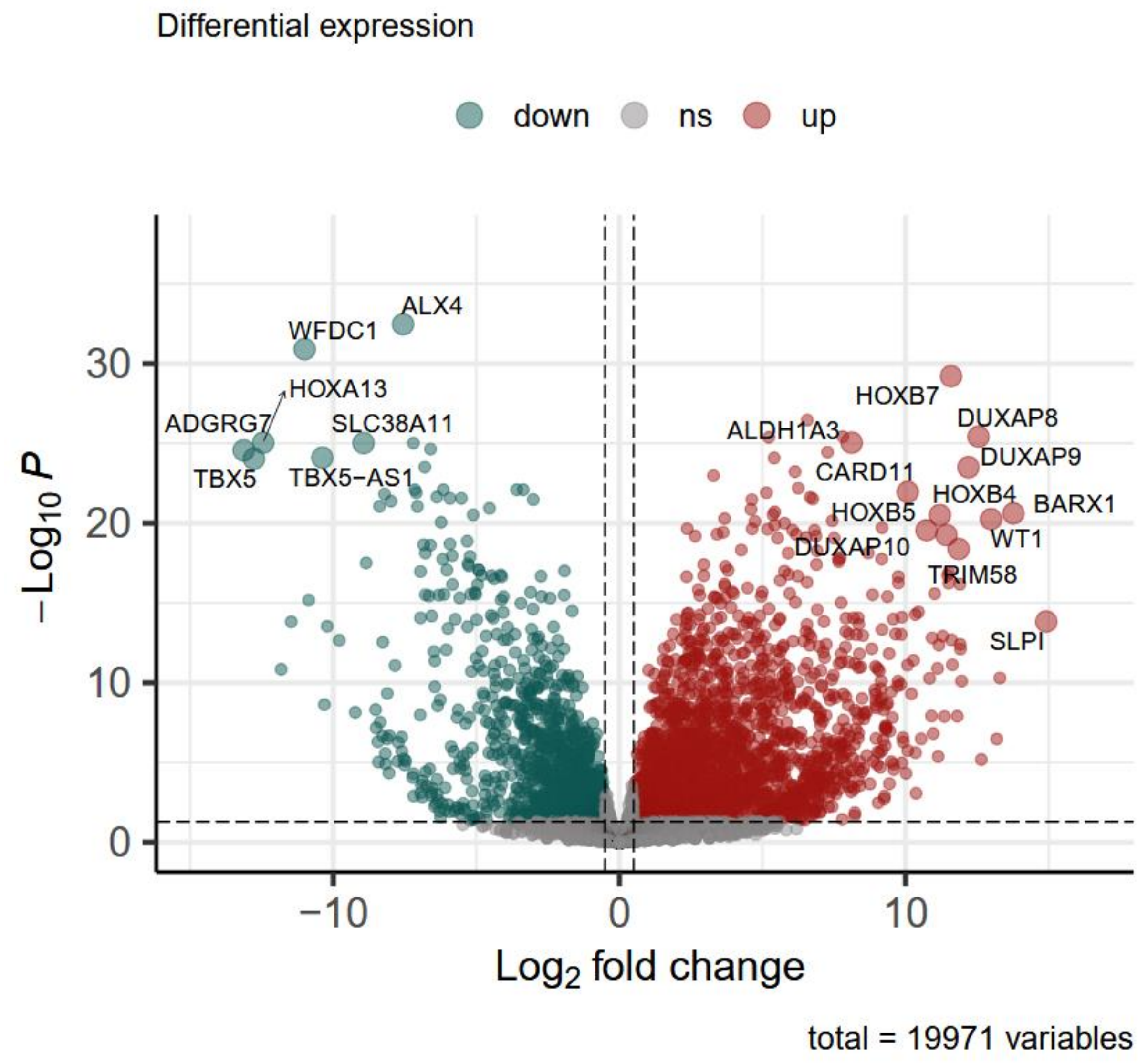


***Figure S9. Stimulus-specific differential gene expression under monopalmitin exposure.*** *Representative volcano plot comparing peritoneal mesothelial cell preparation 5 (P5) following exposure to monopalmitin with matched control conditions. Genes are plotted by $log_2$ fold change (positive values indicate higher expression in monopalmitin-exposed cells; negative values indicate higher expression in controls) and $-log_{10}$ adjusted p-value. Dashed lines denote the thresholds for differential expression ($|log_2FC| \geq 0.5$, FDR < 0.05).*

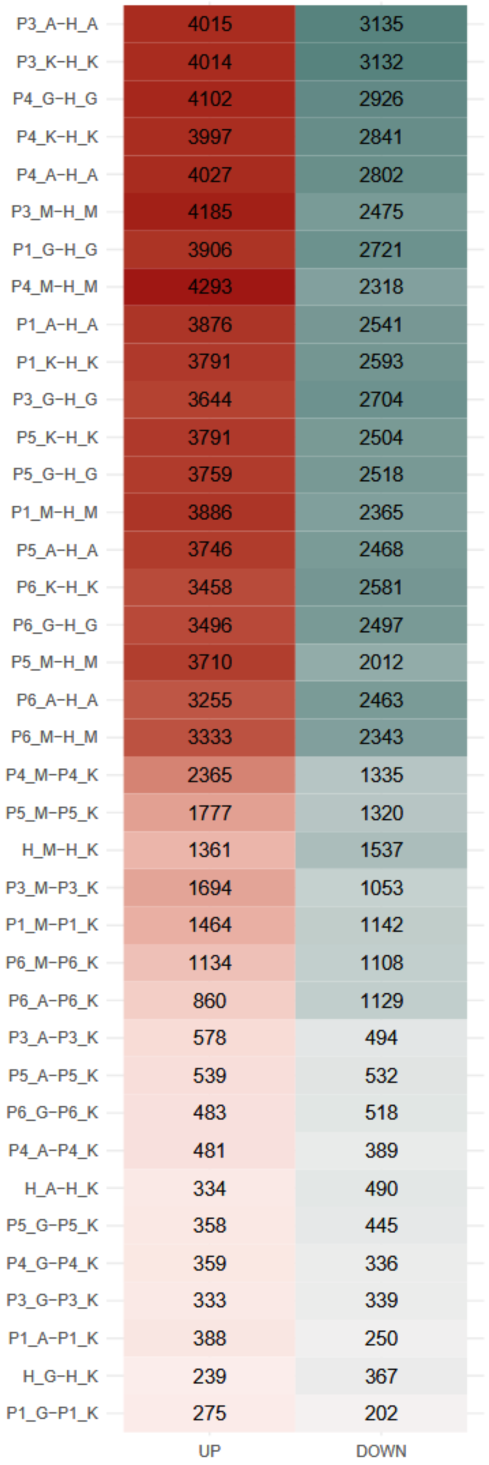


***Figure S10. Quantitative overview of differentially expressed genes across stimuli.*** *Summary of differentially expressed genes (DEGs) showing the number of significantly up- and down-regulated genes per treatment. AES1448 and monopalmitin were associated with a higher number of DEGs compared with GR1479.*

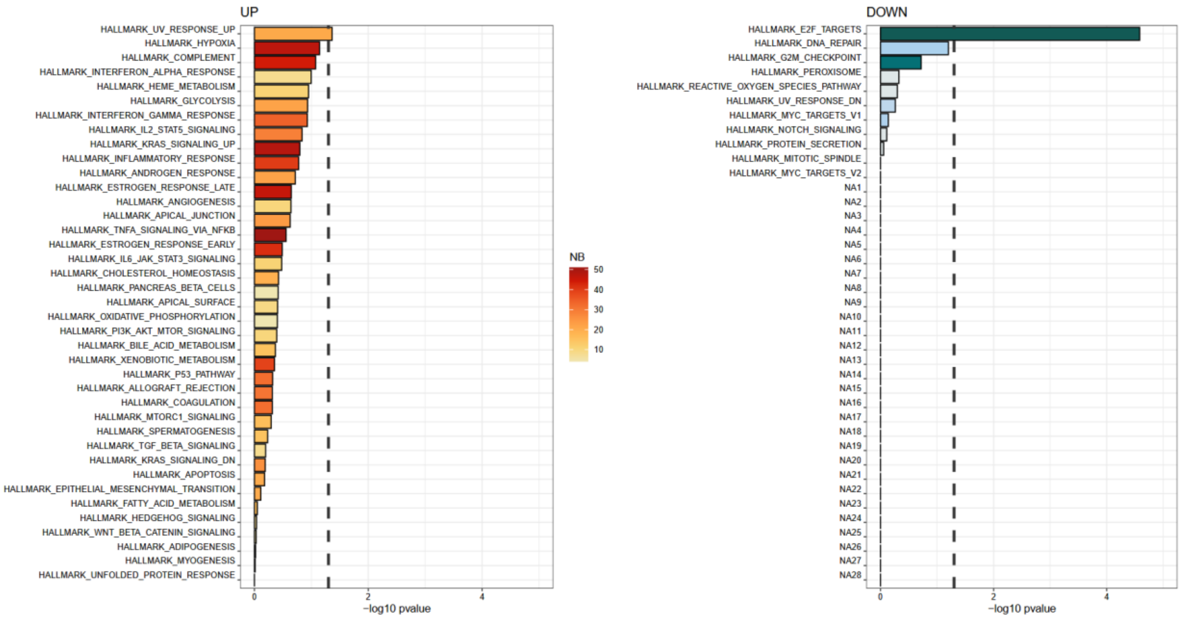


***Figure S11.*** *Condition-specific enrichment profile following GR1479 exposure. Gene set enrichment analysis (GSEA) of peritoneal mesothelial cell preparation 5 illustrates the transcriptional response to GR1479. GR1479 is associated with selective enrichment of UV_RESPONSE_UP and concomitant repression of E2F_TARGETS, indicating a transcriptional shift toward stress-associated programs with reduced proliferative signaling.*

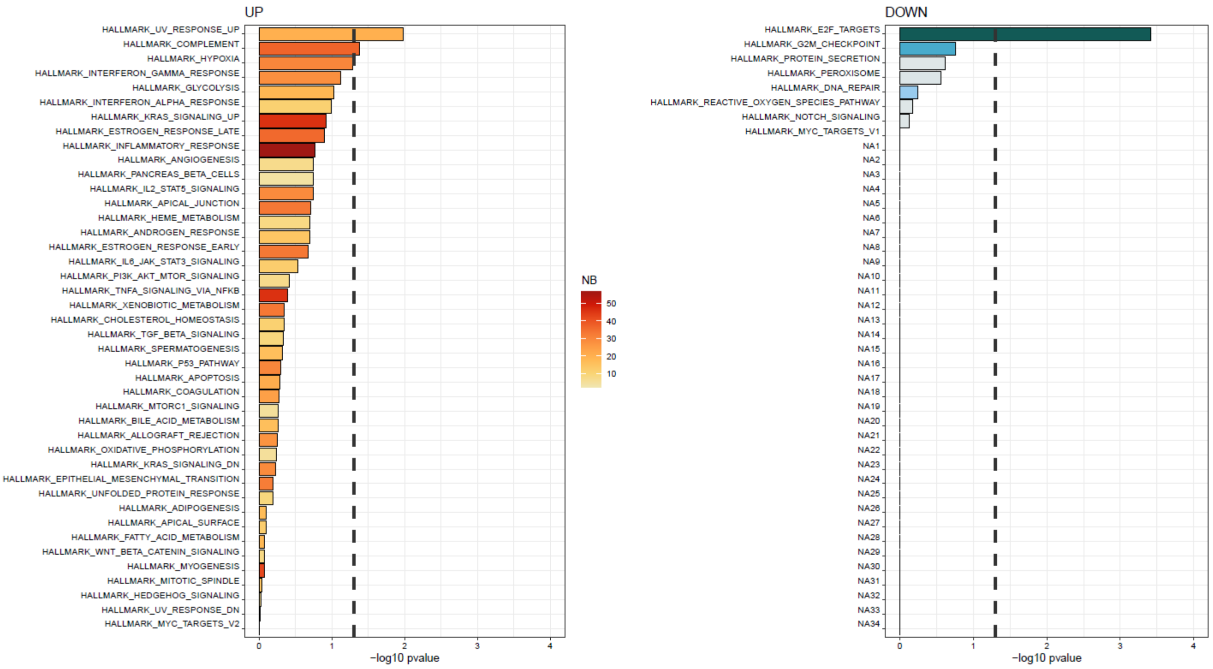


***Figure S12.*** *Condition-specific enrichment profile following monopalmitin exposure. GSEA of peritoneal mesothelial cell preparation 5 demonstrates a distinct enrichment pattern in response to monopalmitin. Pathways related to UV_RESPONSE_UP, COMPLEMENT, and HYPOXIA are enriched, while*

*E2F_TARGETS and G2M_CHECKPOINT are repressed, consistent with a hypoxic–inflammatory stress response accompanied by attenuation of cell-cycle–associated programs.*

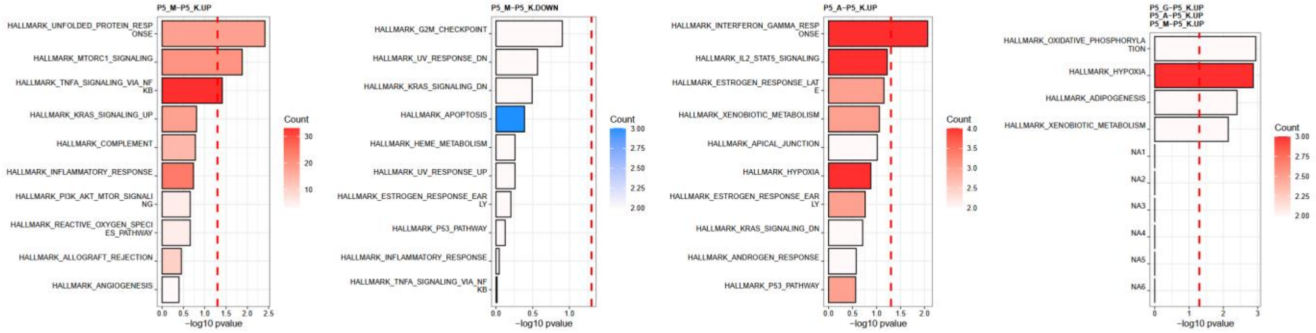


***Figure S13. Extended Fisher and Gene Ontology (GO) biological process analyses.*** *Representative analyses from peritoneal mesothelial cell preparation 5 illustrate stimulus-specific functional enrichment patterns. Monopalmitin is associated with enrichment of acute stress and inflammatory programs, including Unfolded Protein Response and TNFα signaling via NFκB. AES1448 shows predominant activation of immune-related pathways, including Interferon γ Response, IL2–STAT5 signaling, and Hypoxia. In contrast, GR1479 is linked to pathways related to metabolic adaptation, including Oxidative Phosphorylation, Hypoxia, Adipogenesis, and Xenobiotic Metabolism. Together, these profiles delineate two principal axes of cellular response: a toxic–inflammatory axis (NFκB/IFNγ/UPR) and a metabolic–adaptive axis (OXPHOS/Adipogenesis).*

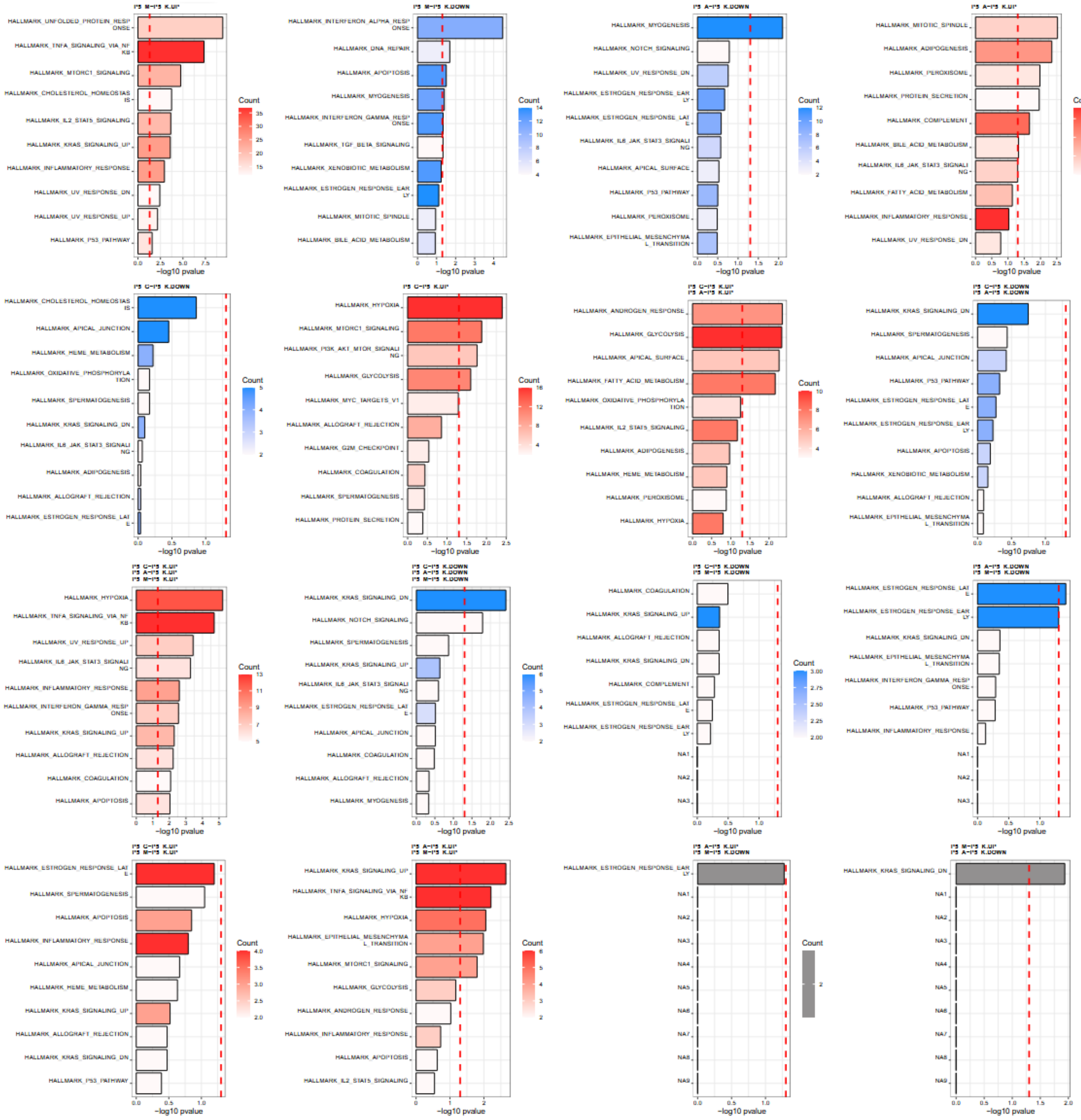

***Figure S14. Hallmark top 10 enrichment overview under relaxed significance thresholds.*** *Robustness analysis based on a representative peritoneal mesothelial cell preparation (P5). Despite the application of relaxed significance thresholds, the overall enrichment patterns remain consistent across conditions. Monopalmitin and AES1448 retain enrichment of stress- and inflammation-associated hallmark pathways, whereas GR1479 continues to show predominance of pathways linked to metabolic adaptation.*

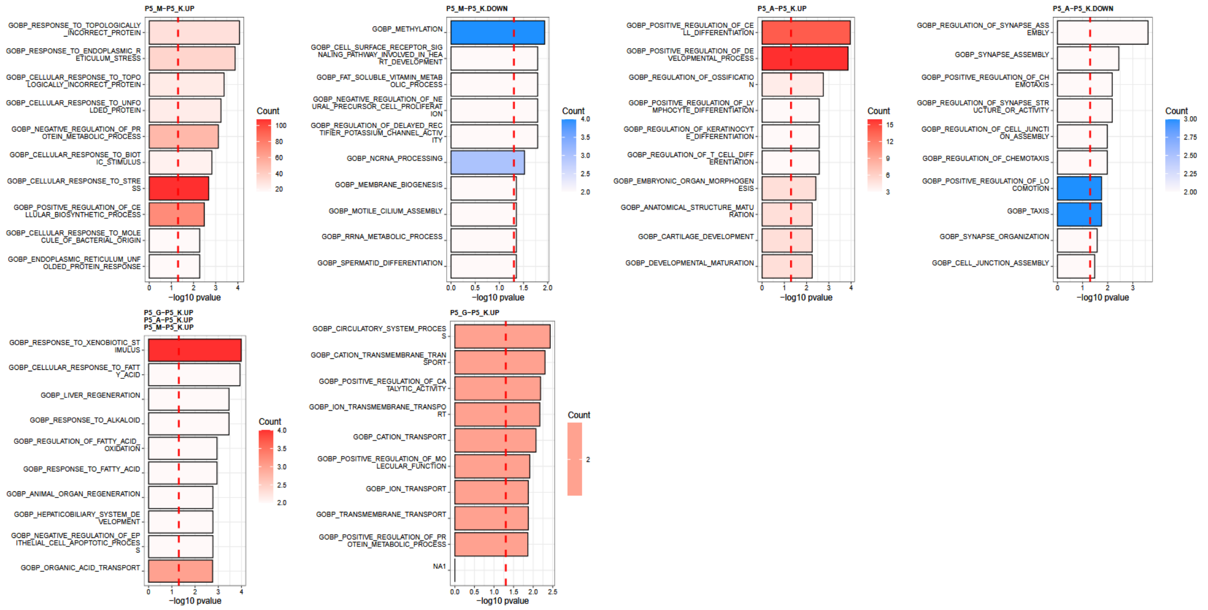


***Figure S15. Gene Ontology (GO) biological process enrichment analysis.*** *Representative analysis from peritoneal mesothelial cell preparation 5 illustrating stimulus-specific biological process enrichment patterns. Monopalmitin is associated with enrichment of processes related to endoplasmic reticulum stress, the unfolded protein response, and general cellular stress responses, accompanied by depletion of biosynthetic processes. AES1448 shows enrichment of terms related to positive regulation of cell differentiation and ossification, together with reduced representation of processes linked to chemotaxis and cell–cell junction organization. In contrast, GR1479 is characterized by enrichment of pathways involved in response to xenobiotic stimulus, fatty acid oxidation, and organic acid transport, consistent with a metabolic adaptive response*

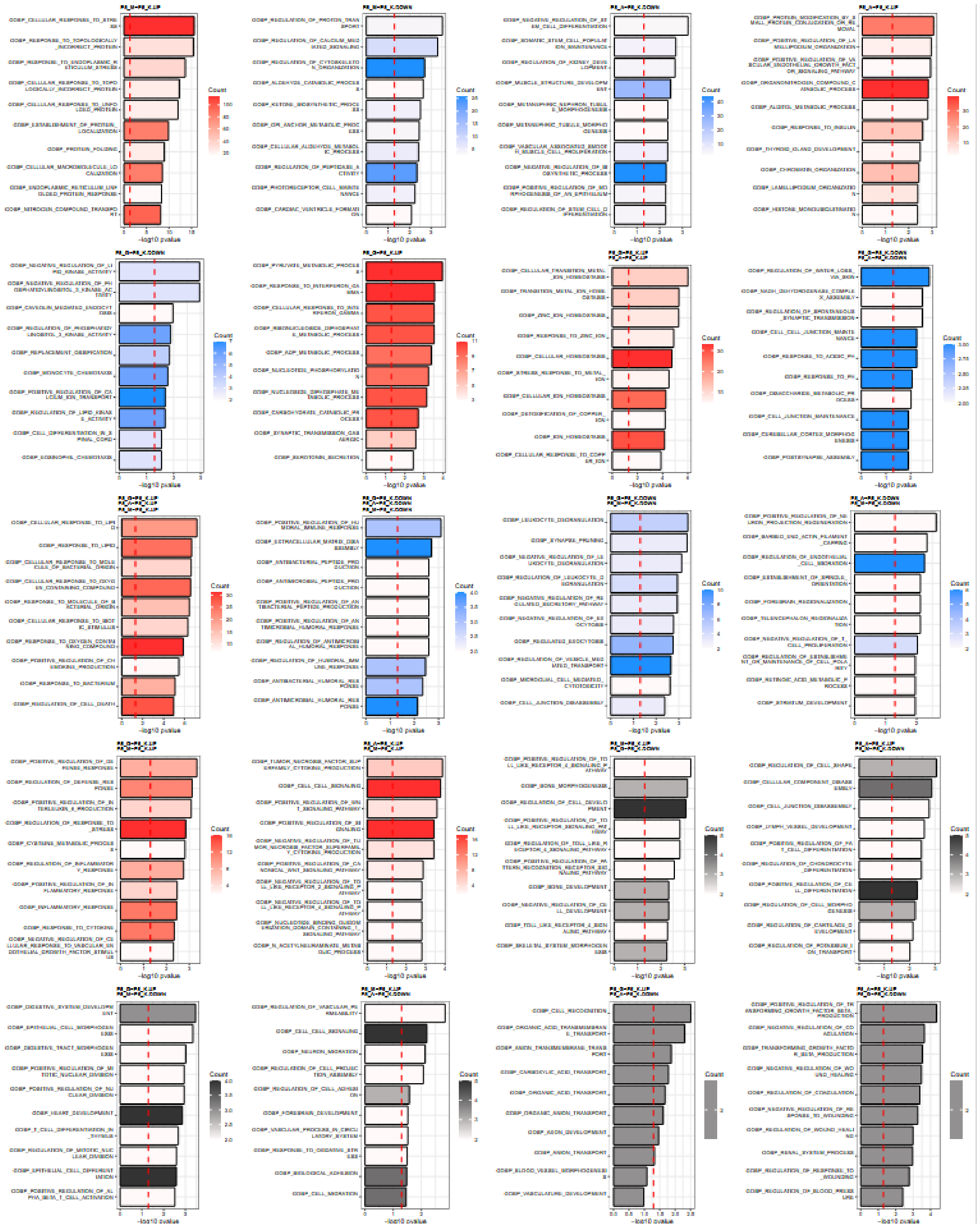

***Figure S16. GO biological process robustness analysis under relaxed significance thresholds.*** *Representative analysis from peritoneal mesothelial cell preparation 5 assessing the stability of GO enrichment profiles. Upon application of relaxed significance thresholds, the overall enrichment patterns remain consistent. AES1448 and monopalmitin retain enrichment of stress- and inflammation-associated biological processes, whereas GR1479 continues to show enrichment of pathways related to metabolic and lipid-regulatory adaptation.*

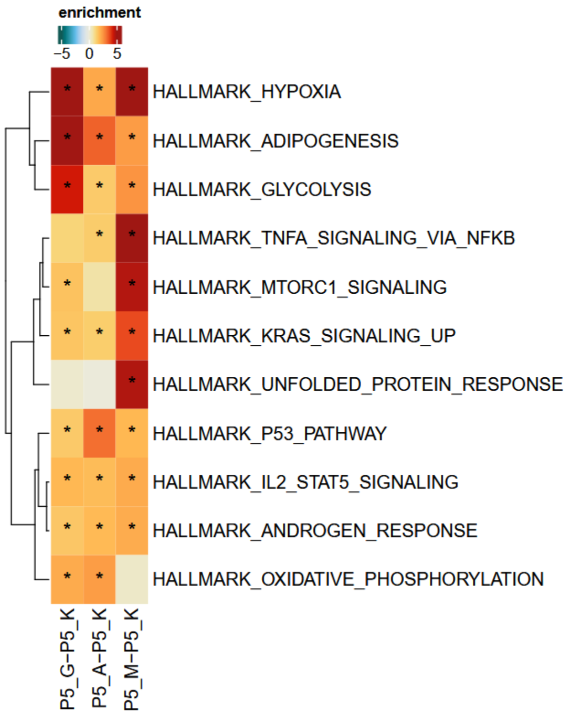


***Figure S17. GSEA heatmap illustrating hallmark pathway enrichment across stimuli.*** *Representative analysis from peritoneal mesothelial cell preparation 5 showing hallmark gene set enrichment following exposure to monopalmitin, AES1448, and GR1479 compared with control conditions. The color scale represents normalized enrichment scores (NES; −5 to +5), and asterisks indicate pathways reaching statistical significance (FDR < 0.05). Pathways related to hypoxia, glycolysis, TNF signaling, and mTORC1 signaling show consistent enrichment across conditions, indicating shared stress-associated transcriptional responses.*
